# Friction patterns guide actin network contraction

**DOI:** 10.1101/2022.12.21.521384

**Authors:** Alexandra Colin, Magali Orhant-Prioux, Christophe Guérin, Mariya Savinov, Ilaria Scarfone, Aurelien Roux, Enrique M. De La Cruz, Alex Mogilner, Manuel Théry, Laurent Blanchoin

**Affiliations:** Univ. Grenoble-Alpes, CEA, CNRS, UMR5168, Interdisciplinary Research Institute of Grenoble, CytoMorpho Lab, 17 rue des Martyrs, 38054 Grenoble, France; Courant Institute of Mathematical Sciences, New York University, New York, NY 10012, United States; Department of Biochemistry, University of Geneva, Geneva, Switzerland; Department of Molecular Biophysics and Biochemistry, Yale University, 260 Whitney Avenue, New Haven, CT 06520-8114, United States; Univ. Paris, INSERM, CEA, UMRS1160, Institut de Recherche Saint Louis, CytoMorpho Lab, Hôpital Saint Louis, 10 Avenue Claude Vellefaux, 75010 Paris, France

## Abstract

The shape of cells is the outcome of the balance of inner forces produced by the actomyosin network and the resistive forces produced by cell adhesion to their environment. The specific contributions of contractile, anchoring and friction forces to network deformation rate and orientation are difficult to disentangle in living cells where they influence each other. Here, we reconstituted contractile acto-myosin networks *in vitro* to study specifically the role of the friction forces between the network and its anchoring substrate. To modulate the magnitude and spatial distribution of friction forces, we micropatterned actin nucleation promoting factors on glass or on a lipid bilayer. We adapted their concentrations on each surface to induce the assembly of actin networks of similar densities, and compare the deformation of the network toward the centroid of the pattern shape upon myosin-induced contraction. We found that actin network deformation was faster and more coordinated on lipid bilayers than on glass, showing the resistance of friction to network contraction. To further study the role of the spatial distribution of these friction forces, we designed heterogeneous micropatterns made of glass and lipids. The deformation upon contraction was no longer symmetric but biased toward the region of higher friction. Furthermore, we showed that the pattern of friction could robustly drive network contraction and dominate the contribution of asymmetric distributions of myosins. Therefore, we demonstrate that during contraction both the active and resistive forces are essential to direct the actin network deformation.

**Significance statement:** Cell shape changes are controlled by complex sets of mechanical forces of various origins. Numerous studies have been dedicated to the role of active forces, originating from molecular motors and filament polymerization, but much less is known about the guiding role of resistive forces. Here we show that a non-uniform distribution of friction forces between a contracting acto-myosin network and its underlying substrate can direct its deformation as it contracts. Our results suggest that the contribution of resistive forces, such as anchoring forces but also less specific viscous forces along cell surface, can be as significant as those of active forces in driving network deformation and should be considered in mechanical models describing the regulation of cell shape and movements.

## Introduction

Actin and myosins are found in all eukaryotes (1–3). Myosins cross-link actin filaments, slide them along each other and thus power large-scale network contraction (4, 5). The regulation of acto-myosin contraction controls cells shape and the mechanical forces they produce on their environment (6–8). In particular, the asymmetry of contraction directs cell deformation and orient tissue morphogenesis (9). Network deformation results from the balance between active contraction and passive resistance. The active contraction depends on the architecture of the actin network and on the amount and spatial distribution of myosins in the network (7, 10). The passive resistance is produced by network anchorages and the friction forces (11, 12). The balance of these four contributions set the rate and orientation of network deformation. Although numerous studies have focused on the mechanism that orient active contraction, the directing role of passive resistance has been much less studied. Can asymmetric friction direct a contractile network as a lagging paddle can guide a canoe?

The **architecture of the actin network** is defined by filament length, density, crosslinking and their respective orientations. It can take the form of branched meshwork or bundles of filaments. Each of these modules have specific contractile properties (13). In cells, their relative dispositions form structural patterns that direct cell deformation and motion (14, 15). Structural anisotropies, such as local weakness induced by local network disassembly, generate the propagation and oscillation of contractile waves (16, 17), the maintenance of which direct the deformation and motion of poorly adhesive cells (18–20). Similarly, local structural reinforcement, by network polymerization in response to the opto-activation of Rac for example, direct cell deformation in the opposite direction (21, 22).

T**he spatial distribution of myosins** depends on their recruitment on actin filaments and activation by signaling pathways (23, 24). Their subcellular distributions generate patterns of contraction, asymmetric cell deformation and migration (25–27). Reconstituted systems were used to show that spatial patterns of myosin activation determines the shape and boundaries of the contracting region of the network (10). In adherent cells, the actin retrograde flow accumulate myosins at the rear of the lamella where they form a subcellular pattern of contractile bundles propelling cells (20, 28–30) and neuronal growth cones (31). In poorly-adhesive cells, a global increase of cell contraction triggers a wave that concentrates myosin in the back and propel cell motion forward (32, 33). Reversal of this contraction pattern is necessary when immune cells reverse their polarity to process the antigens they encounter in the front (34). Sub-cellular activation patterns of myosins could also be induced with opto-genetic tool to induce local contraction and asymmetric deformation contributing to cell motion (20, 35–37). At larger scales, in tissues, asymmetric localization or gradients of myosins power anisotropic tissue deformation (38–41).

**The anchorages** link the actin network to the extra-cellular matrix or to surrounding cells via integrins or cadherins (9). In the local absence of anchorages in the plasma membrane, cortical actin filaments can glide and coalesce to form contractile bundles that concentrate forces (42, 43). By contrast, filaments have limited translocation freedom in regions where anchorages are present so they assemble thinner and pinpointed bundles (42, 44). Thus, extracellular patterns of adhesion are converted into a pattern of structure (45) and a pattern of forces (46). As a consequence, cells tend to detach from regions of lower adhesiveness and move toward regions of higher adhesiveness (47, 48). Such polarization of acto-myosin network deformation can be externally induced by inducing a local detachment of the cell, which is sufficient to induce cell motion toward the opposite direction (49).

**Friction forces** resist the relative sliding of actin network along its anchoring surface (50, 51) and thereby allow cells translocation (52, 53). They also limit the sliding of actin filaments along each other, and thus control network deformation and rearrangements as it contracts (36, 54). In tissues, friction has been proposed to limit the instabilities and direct the self-organization of contractile waves into regular banding patterns (55). This process of directed contraction, in principle, orients network contraction toward regions of higher friction. Although the global role of friction forces has been clearly identified in the migration of poorly or non-adhesive cells (56, 57), or in the motion of encapsulated cell extracts (58), it is still unclear whether pattern of friction can orient cell and tissue deformation. Single cells and cellular aggregates move toward less deformable regions where friction forces between the contracting actin network and the substrate are higher (59–61). At larger scales, anisotropic pattern of friction have been proposed to account for the directed contraction of drosophila apical surface during gastrulation (62), the morphogenesis of the neurectoderm in zebrafish embryos (63), and the emergence of chiral flows in the cortex of C elegans embryos (64). However, in cells, and a fortiori in tissues, it is challenging to distinguish the effect of friction pattern from those of structure, myosin and adhesion patterns. These parameters influence each other and self-organize together (26, 30, 31, 65). Testing the specific role of friction pattern requires to modulate the spatial distribution of friction while all other contributions, structure, myosin and anchorage, are homogeneous. Here we used geometrically-controlled reconstituted acto-myosin networks (66), contracting either on glass or on lipid-coated surfaces (51, 67, 68), to investigate the specific role of friction in the guidance of large-scale contractions.

## Results

### Actin assembly rates are distinct on glass and lipid micropatterns

To study the role of friction on the contraction of acto-myosin networks, we used surface micropatterning to control the initial shape of the network. Our protocol to micropattern actin networks on supported lipid bilayer was inspired from several preceding methods. In brief, glass coverslips coated with a PEG layer were exposed to deep UV through a photomask in order to remove the PEG locally (69). The coverslips were then immersed in a solution containing small unilamelar vesicles, containing 0.25% of lipids conjugated with biotin. The vesicles exploded and spread in contact with the regions presenting bare glass (70). A Nucleation Promoting Factor (Snap-Streptavidin-WA-His) was then attached to these patterned lipids via a biotin-streptavidin link (71). The actin polymerization mix was then added onto this micropatterned surface to generate actin networks of controlled geometries (66) (Figure 1A).

**Figure 1.**
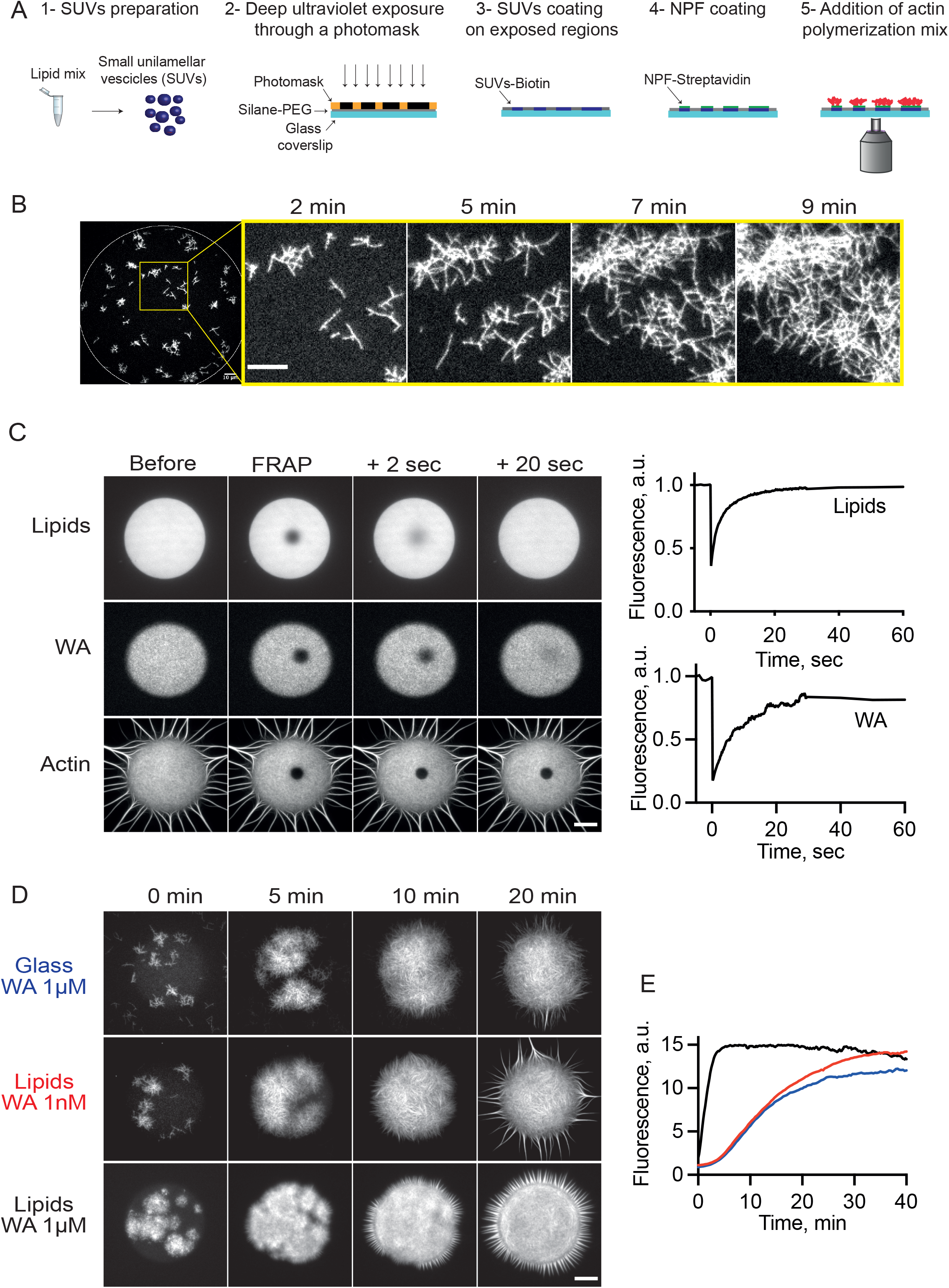
Actin assembly on glass or lipid micropatterns. **A**. Cartoon of the method to constrain branched actin network assembly on glass or lipid micropatterns. **B**. TIRF Imaging of branched actin assembly on lipid micropattern (Disc, diameter 135μm). Biochemical conditions: WA 1nM, Actin 1μM, Human Profilin 3μM, Arp 25nM. Scale bar: 10μm. **C**. Characterization of the diffusion property of the lipid micropattern. Left: TIRF imaging before and after FRAP (zone diameter: 10μm) on lipids, NPF (WA) and Actin (after network polymerization). Biochemical conditions: Disc micropattern (diameter 68 μm), WA 1nM, Actin 1μM, Human Profilin 3μM, Arp 25nM. Scale bar is 20μm. Right: Fluorescence measurements from the images on the left demonstrate that the lipids and the NPF diffuse freely in our experimental conditions. **D**. Comparison of the efficiency of actin network growth on lipid versus glass micropatterns. TIRF imaging of branched actin assembly on lipid and glass disc micropatterns (diameter 68 μm). Biochemical conditions: NPF concentration is indicated on the figure. Actin 1 μM, Human Profilin 3μM, Arp 25nM. Scale bar is 20μm. **E**. Kinetics of actin assembly on lipid versus glass micropattern.

We first checked that NPF was active in our conditions and could induce the assembly of branched actin network on the micropatterned lipid bilayer (Figure 1B, Movie 1), as previously shown on infinite bilayers (72, 73). We also confirmed that lipids and NPF could diffuse freely at normal rates (74) despite the immobility of the dense actin mesh (Figure 1C, Movie 2). Interestingly, we observed that the efficiency to generate an actin assembly with NPF grafted onto a lipid micropattern was an order of magnitude higher (10 times faster) than that of NPF grafted onto a glass micropattern (Figure 1D-E, Movie 3). Considering that network density limits network contraction (73, 75), we determined the concentrations of NPF that induced similar actin network assembly on the glass and lipid micropattern (Figure 1D-E, Movie 3) to investigate specifically the role of substrate friction on network contraction. Hence, in the following experiments the concentration of NPF (Snap-Streptavidin-WA-His) will be 1μM on glass and 1nM on lipids.

### Friction slows down network deformation upon contraction

We assessed the impact of the strength of the interaction between actin nucleation points and the underlying substrate during the actomyosin network contraction. Addition of a double headed (heavy-meromyosin (HMM)-like) myosin VI (13, 75) to a disc-or a square-shaped network on glass or lipid induced its contraction toward the center of the initial shape (Figure 2A-B, Movie 4 and 5).

**Figure 2.**
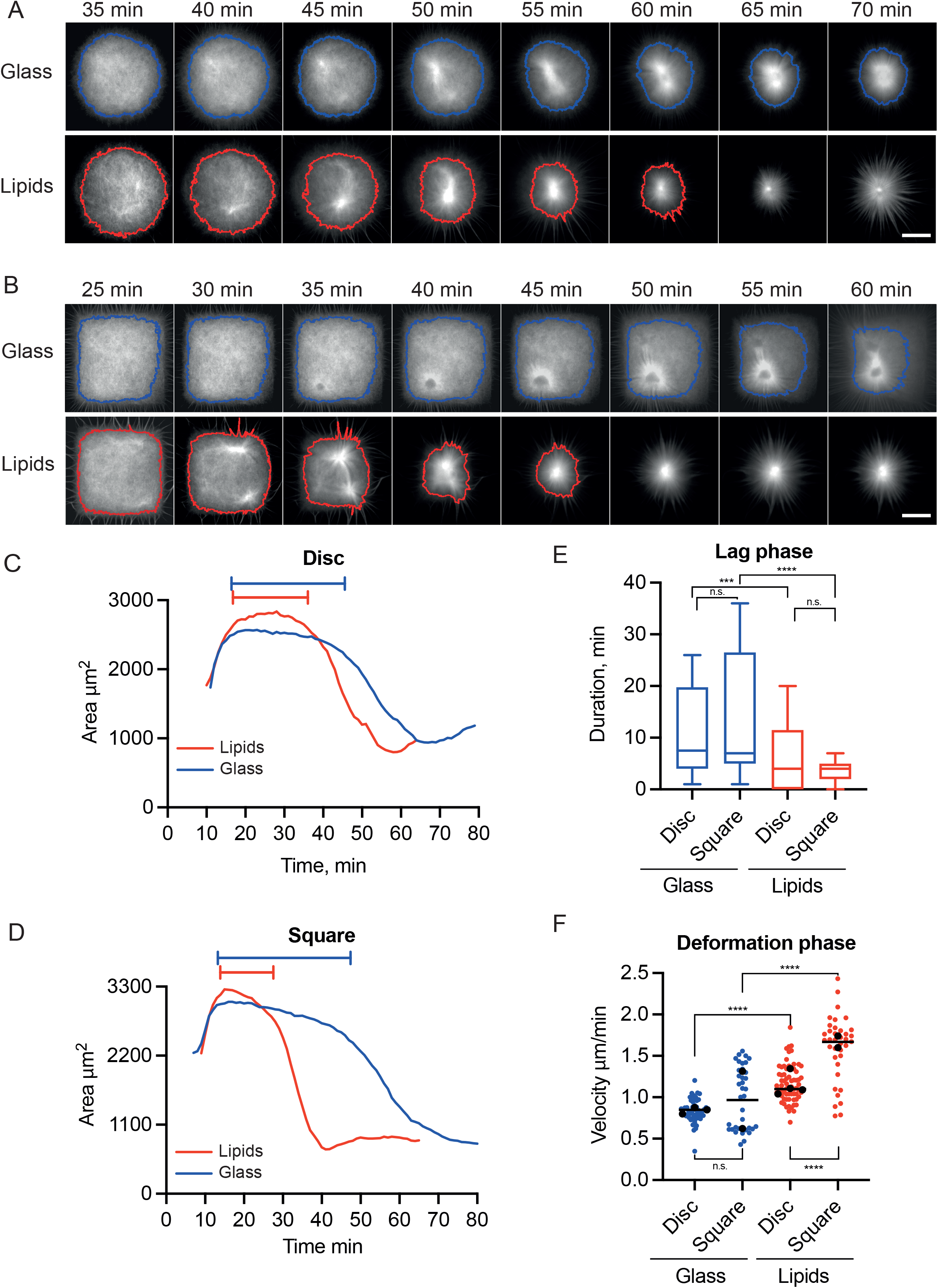
NPFs attachment conditions control actin networks contractile response. **A**. Kinetics of contraction of a disk-shaped actin network on glass or lipid micropattern. Time-lapse imaging of actin network contraction on a glass or lipid disc (diameter 68 μm) micropattern. Line (Blue for glass, Red for lipids) correspond to the contours of the actin network (see methods). Biochemical conditions: On glass micropattern, WA = 1 μM; on lipid micropattern: WA = 1 nM. Actin 1μM, Human Profilin 3μM, Arp 25nM, Myosin VI 14nM. Scale Bar is 20 μm. **B**. Kinetics of contraction of a square-shaped actin network on glass or lipid micropattern. Time-lapse imaging of actin network contraction on a glass or lipid square (length 60 μm) micropattern. Line (Blue for glass, Red for lipids) correspond to the contours of the actin network (see methods). Biochemical conditions: On glass micropattern, WA = 1 μM; on lipid micropattern: WA = 1 nM. Actin 1μM, Human Profilin 3μM, Arp 25nM, Myosin VI 14nM. Scale Bar is 20 μm. **C, D**. Measured actin area as a function of time for the lipid (red) or glass (red) conditions on disc (C) or square (D) micropattern. The period defined at the top of the graph determines the lag phase for each condition. **E**. Duration of the lag phase preceding the contraction for the lipid or glass conditions on disc or square micropattern. Data are represented with boxplots. Disc Glass: n=40-N=2-*median*=7.5. Disc Lipids n=41-N=3-*median* =4.0. Square Glass n=37-N=2-*median* =7.0. Square Lipids n=37-N=2-*median* =4.0. Mann-Whitney Statistics: Disc Glass/Disc Lipids *p* value=0.0010 ***, Square Glass/Square Lipids *p* value ≤0.0001 ****, Disc Glass/Square Glass *p* value=0.1247 ns, Disc Lipids/Square Lipids *p* value=0.5360 ns. **F**. Velocity of the phase contraction phase for the lipid or glass conditions on disc or square micropattern. Data are represented with a superplot. Disc Glass n=53-N=3-*median* =0.8346, Disc Lipids n=68-N=4-*median* =1.133, Square Glass n=37-N=2-*median* =1.013, Square Lipids n=37-N=2-*median* =1.694. Mann-Whitney Statistics: Disc Glass/Disc Lipids *p* value≤0.0001 ****, Square Glass/Square Lipids *p* value≤0.0001 ****, Disc Glass/Square Glass *p* value=0.2761 ns, Disc Lipids/Square Lipids *p* value≤0.0001 ****.

The contraction process followed three phases illustrated by the variation of the area covered by the actin network overtime (Figure 2): An initial growth phase corresponding to the assembly of actin filaments in the first ten minutes after the addition of actin monomer to the reaction mixture; a lag, which is likely corresponding to the loading of myosins on the actin filaments and their local reorganization without any global deformation of the entire network; and the deformation by contraction of the network towards the center of the micropattern (Figure 2C-D). The initial delay of assembly was identical for lipid (red curve) or glass (blue curve) on discs and squares (Figure 2C-D). However, the lag and deformation phases were different on discs and squares. The lag phase was shorter and the deformation rate was faster on lipids than on glass (Figure 2E-F). Similar observations were made for the contraction of networks on rectangles of identical area (Supporting Figure 1). This showed that the lower friction on lipids than on glass facilitates actin network contraction by myosins.

### Friction limits large-scale coordination of network contraction

During this study we noticed that network deformation seemed more homogeneous and coordinated on lipids than on glass. To characterize this, we photobleached a grid in the network and monitored its deformation during the contraction process. We found that on glass, local regions could initiate their deformation independently, while on lipids, the deformed regions formed a continuum and contracted together across the network (Figure 3A). To further quantify this, we tracked actin network deformation by particle image velocimetry (Figure 3B and C). For the analysis we decomposed the square in four quadrants and summed the velocities of the moving parts in each quadrant. Measurements showed that the deformation of the four quadrants were not similar nor synchronized on glass but they were precisely coordinated on lipids (Figure 3B-C and Supporting Figure 2). Interestingly, this difference of spatial and temporal coordination during contraction had an impact on the position of the point of convergence of the contraction process, which was closest to the center of the initial shape on lipids than on glass (Figure 3D, Movies 6 and 7). This showed that the lower friction on lipids than on glass allowed a more integrated and coordinated acto-myosin contraction.

**Figure 3.**
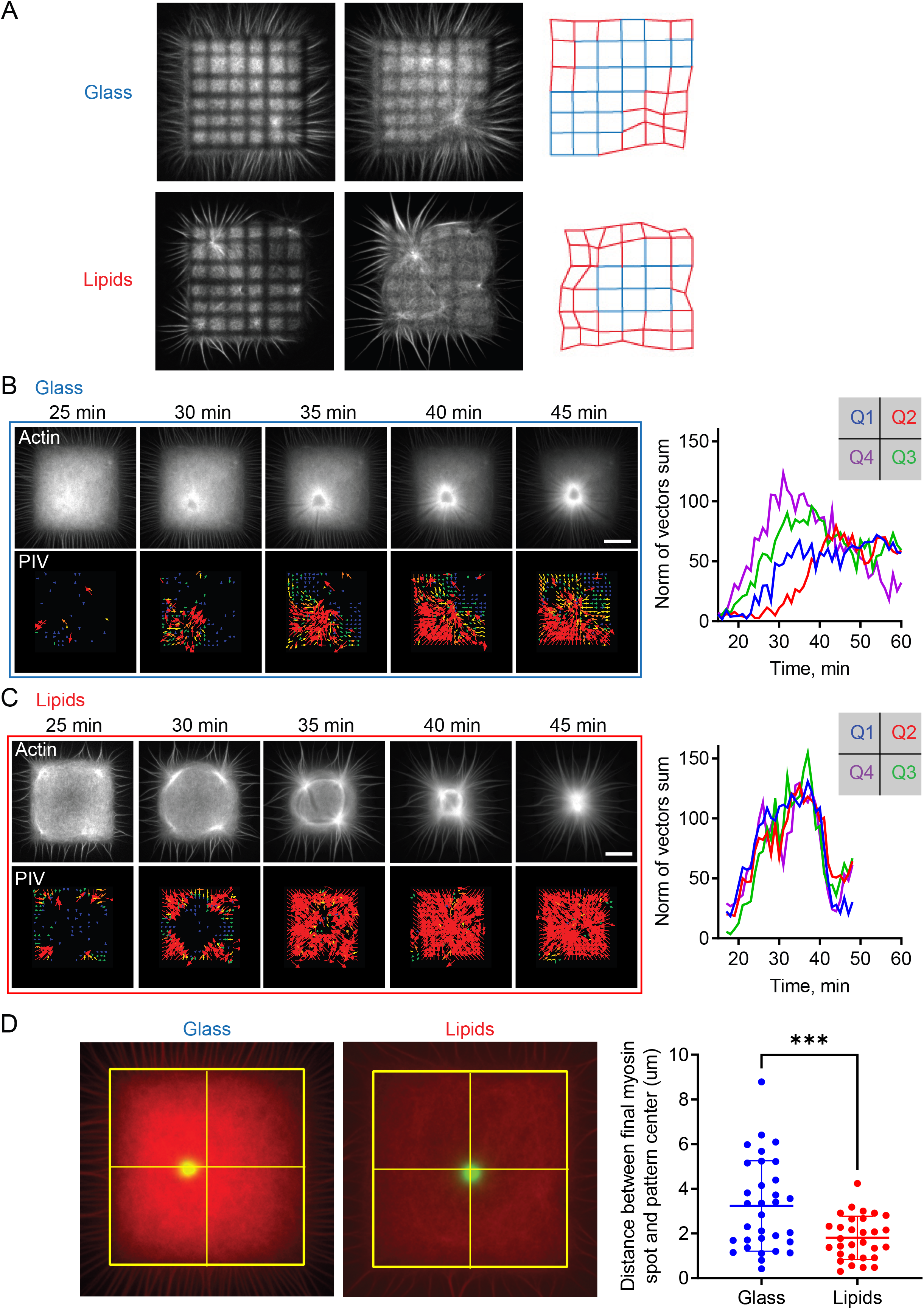
Lower friction improves the coordination of the contraction process. **A**. Local deformation of actin network. A photobleached grid shape was performed on actin networks polymerized on glass or lipid micropatterns. Then deformations of the actin network were followed by following the grid deformation. **B**. Left Top: Example of an actin network grown on glass substrate pattern used for PIV analysis. Left Bottom: PIV analysis of the actin network shown above. Right: Resultant of vector sum for each quadrant defined on the pattern as a function of time (see Methods). **C**. Same as B with an actin network grown on lipid substrate. **D**. Left: Snapshots of final position of myosin spot for actin networks grown on glass or lipid square micropattern. Right: quantification of the distance between the final myosin spot and the pattern center for networks grown on glass or lipids. N = 2 independent replicates with n = 18 and n = 12 patterns for the lipids condition and n= 19 and n = 13 patterns for the glass condition. Individual points for each pattern are represented. Mean and standard deviation are plotted on top of the points. Unpaired t-test: *p* value≤0.001 ***. Biochemical conditions for Figure 3: On glass micropattern, WA = 1 μM; on lipid micropattern: WA = 1 nM. Actin 1μM, Human Profilin 3μM, Arp 25nM, Myosin VI 14nM.

### A pattern of friction can guide network contraction

The above-mentioned effect of friction on the rate and coordination of the contraction process suggested that heterogeneity in the friction coefficient between the actin network and the underlying substrate might direct network contraction. To test this hypothesis, we developed a new method to generate a heterogeneous friction pattern below uniform and geometrically controlled actin networks (Supporting Figure 3A). We used a digital micromirror device to photo-activate distinct regions sequentially and coat them with distinct components (see Methods and Supporting Figure 3A). On a PEG layer, we generated discs, squares or rectangles, half of which was made of bare glass coated with 1μM NPF and the other half was coated with lipids and 1nM NPF in order to obtain similar actin polymerization kinetics on the two halves and thus a homogeneous network over the entire micropattern. We confirmed that lipids and WA diffused on the lipid half and not on the glass half (Supporting Figure 3B) and that the networks contracted according to the three phases described previously (Supporting Figure 4).

Strikingly, while networks on homogeneous disk-shape lipid micropatterns contracted precisely toward the centroid of the initial geometry, they showed a clear off-centering toward the region of higher friction on heterogeneous glass-lipid micropatterns (Figure 4 A,B, and Movies 9-14). This deviation resulted from the faster deformation of the acto-myosin network on lipids than on glass (Supporting Figure 4). We tested additional patterns of friction in squares and rectangles and noticed that the network compacts to a point which is biased toward the center of the higher friction region (Figure 4C-F). These results demonstrated that a friction pattern can guide acto-myosin network deformation.

**Figure 4.**
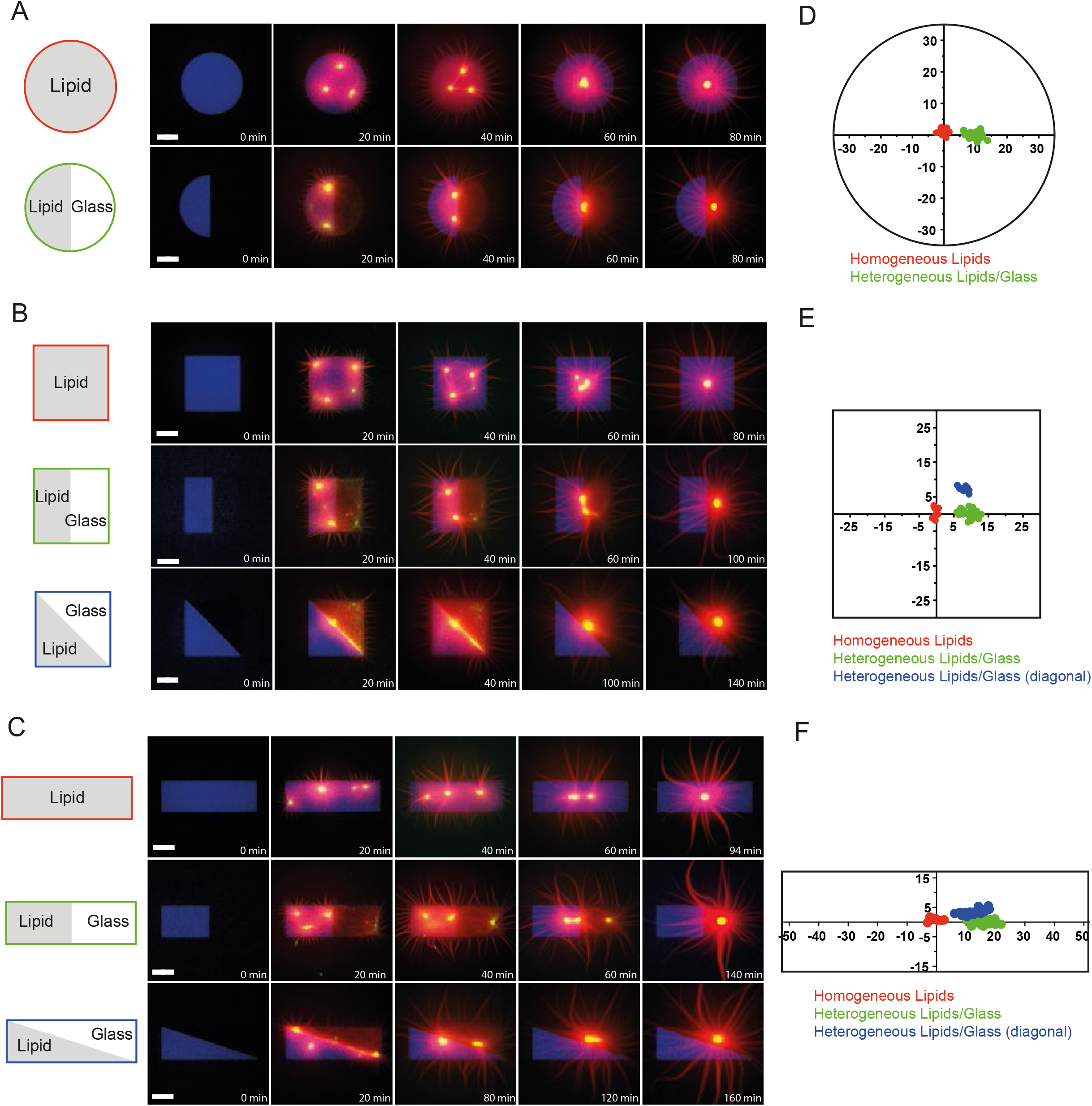
Friction pattern directs network contraction. Time lapse imaging of actin network contraction on full lipid or heterogeneous discs **(A)**, full lipid or heterogeneous squares **(C)**, full lipid or heterogeneous rectangles **(E)**. Biochemical conditions: Actin 1 μM, Human Profilin 3 μM, Arp 25 nM, Myosin VI 10 nM. Patterns dimensions: Discs, 64 μm diameter; Squares 60 μm length; Rectangles 104 μm x 34 μm. Pattern is represented in blue, actin in red, myosin in green. Scale bar: 20 μm. **B, D, F**. Plot of the myosin dots coordinates at the end of contraction for the different shapes. All coordinates were measured and compared to the centroid (x=0, y=0) of the whole pattern. Red dots correspond to the coordinates of the full pattern made of lipid, in green the dot correspond to the coordinates of the symmetrical heterogeneous pattern and the blue dots are the coordinates of the asymmetrical heterogeneous patterns.Disc full lipids: N = 3, n = 11 patterns. Heterogeneous disc: N = 4, n = 26 patterns. Square full lipids: N = 2, n = 9 patterns. Heterogeneous square: N = 4, n = 29 patterns. Asymmetrical heterogeneous square: N = 2, n = 11 patterns. Rectangle full lipids: N = 2, n = 8 patterns. Heterogeneous rectangle: N = 4, n = 24 patterns. Asymmetrical heterogeneous rectangle: N = 3, n = 21 patterns.

### Numerical simulations account for the trajectories of myosin spots during network contraction

To further understand the way acto-myosin networks deform as they contract, we turned to computational modeling. We model the actomyosin network as a two-dimensional viscoelastic network with active contractile stresses generated by myosin concentrated in several foci (76). The internal (passive viscoelastic and active contractile) stresses in the network are balanced by friction between the network and substrate (glass or lipid), which we model as effective viscous drag with the drag on glass higher than that on lipid. This model is implemented as a network of nodes connected by a viscous dashpot with the substrate. Neighboring nodes are connected by the viscous dashpot and elastic (nonlinear) spring in series responsible for viscous and elastic deformations inside the network (see Model description in supplementary information and Figure 5A).

**Figure 5.**
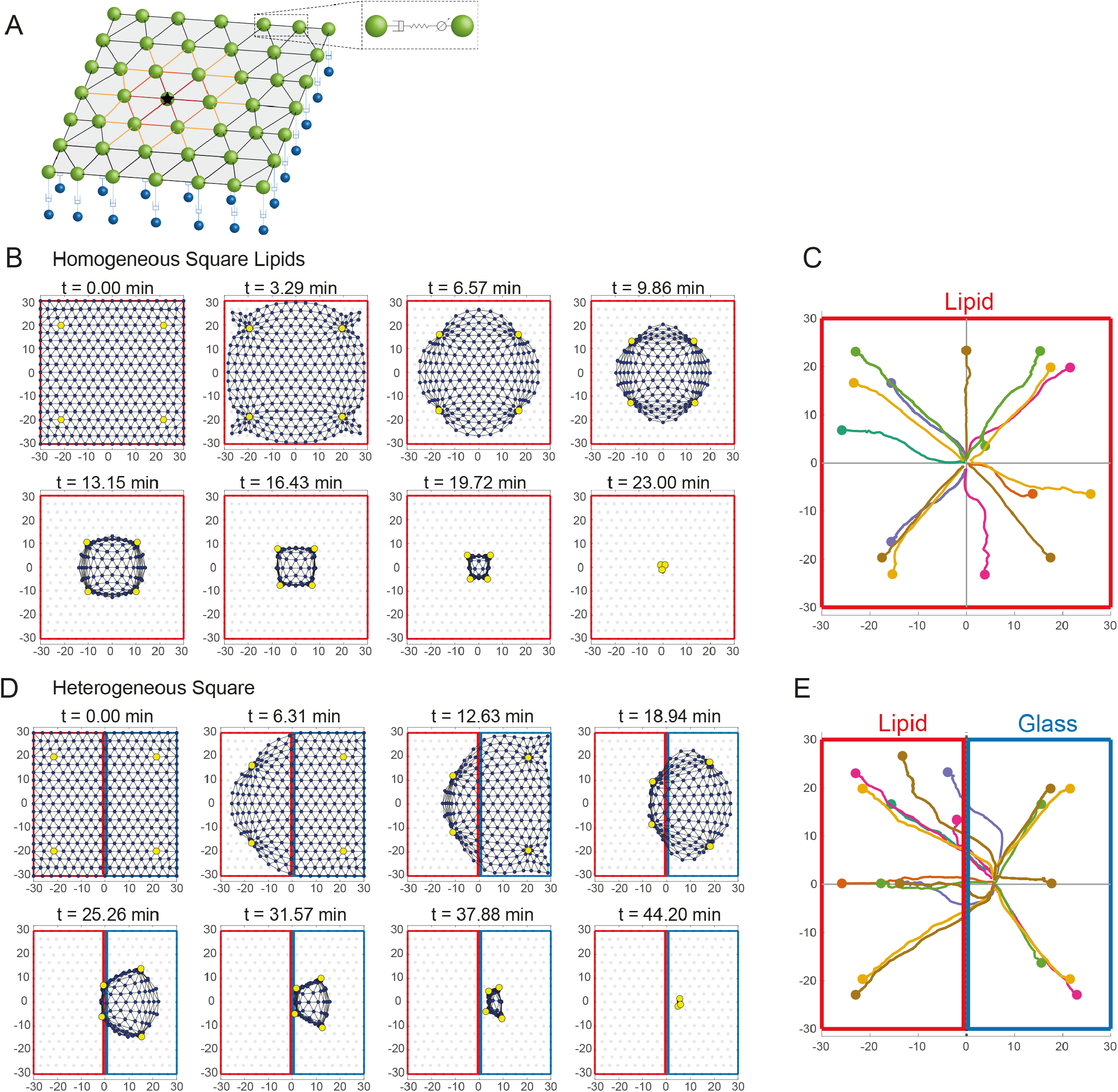
Computational model recapitulates the contraction kinetics and predict the trajectories of myosin spots during the network contraction. **A**. Scheme of the computational model describing the network as a viscoelastic network of nodes and links. Each node is connected by a dashpot with the substrate. The dashpot is responsible for the friction (higher on glass, lower on lipid). Neighboring nodes are connected by dashpot/spring in series responsible for viscous and elastic deformations inside the network. In addition, links connecting pairs of nodes, one of which is occupied by a myosin dot (shown by star), are contractile. **B**. Sequence of deformations predicted by the model for an actin network polymerized on a homogeneous square lipid micropattern and contracted by myosin dots (yellow nodes). **C**. Prediction of myosin trajectories for an actin network contracting on a homogeneous square lipid micropattern. **D**. Sequence of deformations predicted by the model for an actin network polymerized on a heterogeneous square micropattern and contracted by myosin dots (yellow nodes). **E**. Prediction of myosin trajectories for an actin network contracting on a heterogeneous square micropattern.

We first simulated the contraction of the network on homogeneous lipid square, with initial myosin foci appearing in four corners of the square (Figure 5B, Movie 16). The simulations showed the temporal sequence of the network deformation like those observed experimentally (Figure 2B). Specifically, the simulations correctly predict that the network corners get rounded right after the onset of the contraction (Figure 5B). Note also that the area near the center of the network did not deform until late stage of contraction (Figure 5B), which is also observed in experiments (Figure 3A). We then plotted the predicted myosin foci’s trajectories for homogeneous patterns. Interestingly, for the homogeneous pattern, the model predicts that regardless of the initial positions and even number of myosin spots, the spots drifted straight to the center of the square and converged to it (Figure 5C).

Remarkably, when a heterogeneous pattern of friction was simulated with the exact same initial myosin distribution, the final point of contraction was shifted toward the side of high friction as observed on the experimental data (Figure 5D, Movie 17). Furthermore, for the square homogeneous or heterogeneous patterns we found that regardless of the initial position and number of myosin foci, the foci drifted straight to the center of the friction pattern and then converged to it (Figure 5C, D and see Supporting Figure 5A-B for examples with different positions of myosin foci). This is reminiscent of the ballistic contractility previously described (76). Finally, the trajectories predicted on homogeneous or heterogeneous rectangles showed myosin foci first converging to the point of intersection of the bisecting lines of the angles and then moving to their destination, defined by the distinct weights of the two friction patterns. (Figure Supporting Figure 5C-F).

Two contraction features are illustrated by these trajectories: first the global character of the contraction: myosin foci integrate large areas in the definition of their motion, which is largely the consequence of the highly interconnected character of the actomyosin network and global propagation of the internal stresses (see model description in supplemental information). Second, the predicted locations of the convergence points of the contracted heterogeneous pattern are largely insensitive to the initial myosin distributions (and to the number of myosin foci) but determined by the adhesion distribution.

As a consequence, heterogeneity of contractility (initial myosin distribution) does not explain nor impact contraction asymmetry. Although the contraction is local and randomly initialized by myosin foci, a combination of network geometry and external friction heterogeneity leads to a robust global polarized contraction. Thus, our model proposes that, the friction pattern can dominate over the initial distribution of contractility to drive contraction asymmetry.

### Friction acts as a master control of symmetry of actomyosin meshwork contraction

We tested the predictions of the model by analyzing the localization and trajectories of myosin foci in our experimental data. Foci formed rapidly and were randomly distributed in networks assembled on homogeneous square lipid micropatterns (Figure 6A). The heterogeneity of these initial distributions, which could be quite asymmetric, and sometimes even made of a single and off-centered focus, did not impact the position of the convergence point, at the square center, nor the trajectories, which were directed straight toward it (Figure 6A, movie 10). These observations were consistent with the model (Figure 5B-C) and demonstrate that the acto-myosin network contraction integrates the entire shape of the network independently of the asymmetry of myosins distribution.

**Figure 6.**
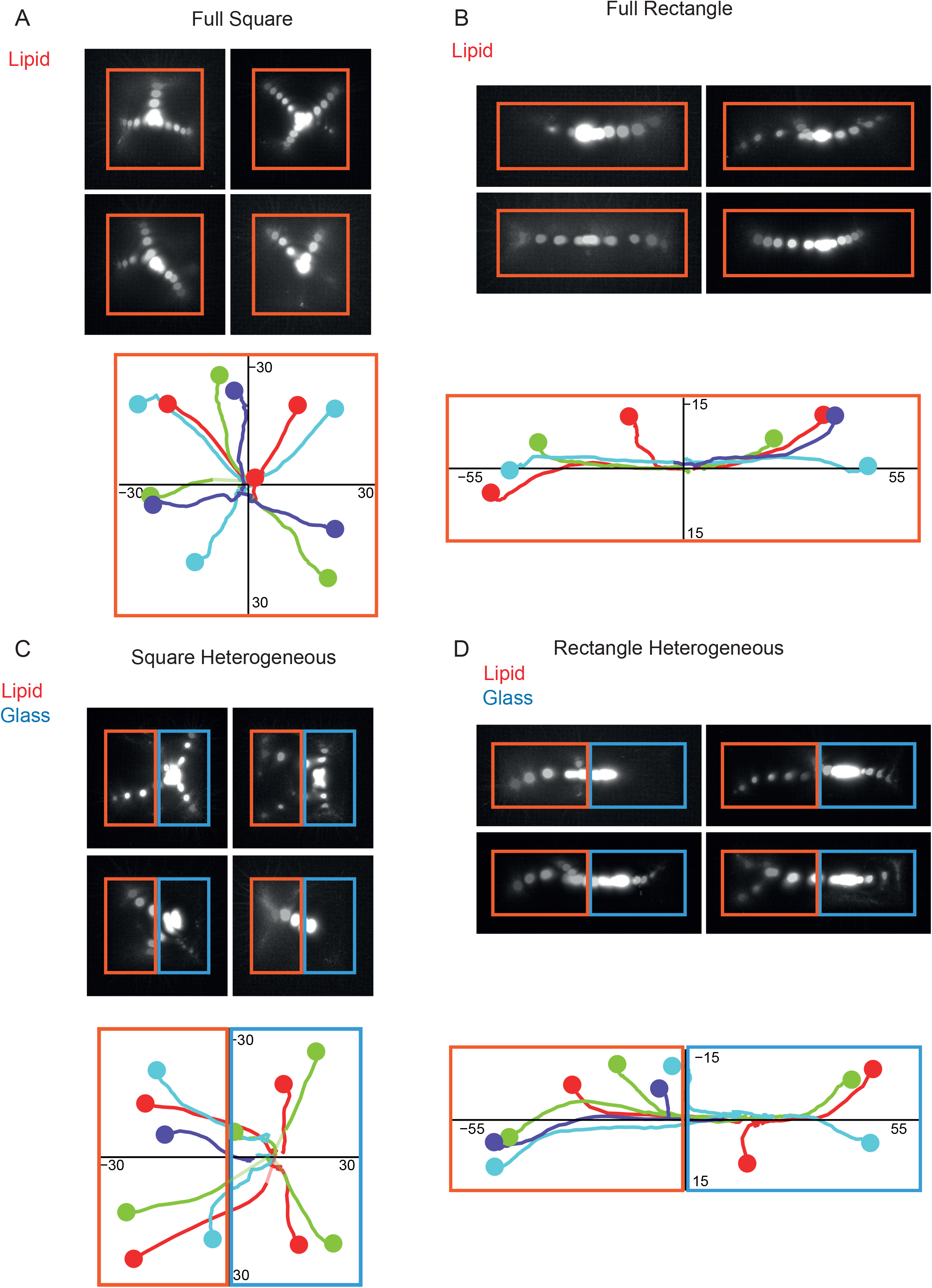
Friction pattern robustly drives network contraction despite uneven distribution of myosin. Top: Four examples of temporal projections of myosin detection on a full-square micropattern **(A)**, a full rectangle micropattern **(B)**, a heterogeneous square micropattern **(C)**, a heterogeneous rectangle micropattern **(D)**. Bottom: Detection of myosin spots as a function of time (see Methods for details). Each example of the above images is represented with a single color. The initial time of the trajectory is represented with a bigger dot.

Tracking myosin foci on homogeneous rectangular lipid micropatterns confirmed the above conclusions on the absence of impact of the asymmetry of myosin distribution. They also showed that myosin foci first moved along angle bisecting lines and then reorient toward the rectangle center. These observations were also consistent with the model (Figure 5C, E and Supporting Figure 5E-F) and further demonstrated that the contraction process integrate local and global network geometry, as previously shown (10), regardless of the initial pattern of myosins.

Importantly, during the contraction of actin networks assembled on heterogeneous square patterns of glass and lipids, we also found that myosin foci could be observed above both types of anchors (Figure 6C, movie 11-12). This confirmed that network density and architectures were similar on glass and on lipids and that the initial localization of myosin foci was not determined by the strength of the underlying anchorage. The trajectories were initially not all directed toward network final convergence point, likely due to the asynchrony of the contraction described earlier (Figure 2) but then all move toward the off-centered destination regardless of their position above glass or lipids, further confirming that the network integrated the entire geometry and the entire heterogeneity of the underlying friction pattern (Figure 6C). Foci trajectories during the contraction of actin networks assembled on heterogeneous rectangular patterns of glass and lipids confirmed these two conclusions on the dominant roles of network geometry and friction pattern over the initial distribution of myosins (Figure 6D).

## Discussion

Actin network architecture, myosin distribution and network anchorages are intimately related. They co-assemble and influence each other in complex feedback loops. Indeed, adhesion primes filament assembly (77, 78), dense filament areas recruit more myosins (79, 80) and contractile forces promote the enlargement of adhesions (81, 82). It is therefore difficult to attribute the deformation process of the contracting network to one or the other parameter. Most of previous works were focused on the steering role of motors distribution since it powers the contraction process (83, 84). In vitro experiments revealed the role of network architecture in modulating the rate of network deformation by its impact on the connectivity of the contractile network (13, 73, 75). Here we clearly establish the key role of friction pattern in the guidance of network deformation during contraction. This implies for cells and tissues that homogeneous contractile networks, made of even distribution of actin filament and myosins, can contract asymmetrically if their anchorages are not evenly distributed, or if the strength of these anchorages are not identical all over the network.

Our numerical simulations of a visco-elastic contractile network and our experimental data also shows the less intuitive result that the friction pattern can guide network contraction in a quite robust manner that dominates the contribution of a heterogeneous distribution of myosins. In physiological conditions, this implies that morphogenetic processes can result from the local recruitment and activation of myosins by signaling but that the role of adhesion is also key in the establishment of the final orientation of the deformation process. Thus, both orientations of the powering force and the resistive forces are essentials, as well known by sailors which efficiently combine the use of wind orientation in the sails to propel the boat and the rudder blade orientation to establish its final direction.

## Materials and methods

### Protein expression, purification, and labeling

Actin was purified from rabbit skeletal-muscle acetone powder (85). Monomeric Ca-ATP-actin was purified by gel-filtration chromatography on Sephacryl S-300 at 4°C in G buffer (2 mM Tris–HCl, pH 8.0, 0.2 mM ATP, 0.1 mM CaCl_2_, 1 mM NaN_3_ and 0.5 mM dithiothreitol (DTT)). Actin was labelled on lysines with Alexa-568 (86). All experiments were carried out with 5% labelled actin. The Arp2/3 complex was purified from calf thymus according to (87) with the following modifications: the calf thymus was first mixed in an extraction buffer (20 mM Tris pH 7.5, 25 mM KCl, 1 mM MgCL2, 0.5 mM EDTA, 5% glycerol, 1 mM DTT, 0.2 mM ATP and proteases). Then, it was placed in a 50% ammonium sulfate solution in order to make the proteins precipitate. The pellet was resuspended in extraction buffer and dialyzed overnight. Human profilin was expressed in BL21 DE3 pLys *Escherichia coli* cells and purified according to (88). Double-headed porcine myosin VI with bound calmodulin was purified from Sf9 cells by FLAG affinity chromatography (89, 90).

Snap-Streptavidin-WA-His (pETplasmid) was expressed in Rosettas 2 (DE3) pLysS (Merck, 71403). Culture was grown in TB medium supplemented with 30 μg/mL kanamycine and 34 μg/mL chloramphenicol, then 0.5 mM isopropyl β-D-1-thiogalactopyranoside (IPTG) was added and protein was expressed overnight at 16 °C. Pelleted cells were resuspended in Lysis buffer (20 mM Tris pH8, 500 mM NaCl, 1 mM EDTA, 15 mM Imidazole, 0,1% TritonX100, 5% Glycerol, 1 mM DTT). Following sonication and centrifugation, the clarified extract was loaded on a Ni Sepharose high performance column (GE Healthcare Life Sciences, ref 17526802). Resin was washed with Wash buffer (20 mM Tris pH8, 500 mM NaCl, 1 mM EDTA, 30 mM Imidazole, 1 mM DTT). Protein was eluted with Elution buffer (20 mM Tris pH8, 500 mM NaCl, 1 mM EDTA, 300 mM Imidazole, 1 mM DTT). Purified protein was dialyzed overnight 4°C with storage buffer (20 mM Tris pH8, 150 mM NaCl, 1 mM EDTA, 1 mM DTT), concentrated with Amicon 3KD (Merck, ref UFC900324) to obtain concentration around 10 μM then centrifuged at 160000 x g for 30 min. Aliquots were flash frozen in liquid nitrogen and stored at −80 °C.

### SUV (small unilamellar vesicules) preparation

L-α-phosphatidylcholine (EggPC) (Avanti, 840051C), DSPE-PEG(2000) Biotin (1,2-distearoyl-sn-glycero-3 phosphoethanolamine-N-[biotinyl(polyethylene glycol)-2000], ammonium salt, Avanti, : 880129C-10mg chloroforme) and ATTO 647N labeled DOPE (ATTO-TEC, AD 647N-161 dehydrated) were used. Lipids were mixed in glass tubes as follows: 98.75% EggPC (10 mg/mL), 0.25% DSPE-PEG(2000) Biotin (10 mg/mL) and 1% DOPE-ATTO390 (1 mg/mL). The mixture was dried with nitrogen gas. The dried lipids were incubated under vacuum overnight. After that, the lipids were hydrated in the SUV buffer (10 mM Tris (pH 7.4), 150 mM NaCl, 2 mM CaCl_2_). The mixture was sonicated on ice for 10 minutes. The mixture was then centrifuged for 10 min at 20,238 x g to remove large structures. The supernatants were collected and stored at 4°C until use.

### Preparation of SilanePEG passivated slides

SilanePEG (30kDa, PSB-2014 or 5 kDa, PLS-2011, Creative PEG works) was prepared at a final concentration of 1 mg/mL in 96% ethanol and 0.1%(v/v) HCl. Slides and coverslips were cleaned with the following protocol: they were sonicated for 30 minutes at 60°C in Hellmanex 2%. They were then rinsed with several volumes of mqH2O. Just before use, they were dried with compressed air. Slides and coverslips were plasma cleaned (Femto; Diener Electronic) for 5 minutes at 80% power and directly immersed in the silanePEG solution for 18 hours. They were dried just before use.

### Lipid micropatterning

Deep UV exposure through a chrome-photomask during 45 seconds creates micropatterns on SilanePEG coverslip. The coverslip is then mounted in a 70 μm height flow chamber (with a silanePEG slide on top, double-sided tape, Lima ref 1820080). 30 μL of SUV solution is then immediately perfused in the flow chamber. After 10 minutes of incubation at room temperature, the flow chamber is rinced with 1 mL of SUV buffer. Diffusivity of the lipid bilayer was checked before each experiment with a FRAP on the lipid bilayer.

### Actin polymerization and contraction

Micropatterning coating (with and without bilayer of lipids) was done with WA protein labeled with snap488 for polymerization experiment and no-labeled WA protein for contraction. WA was incubated for 10 minutes at room temperature on micropatterns. The flow chamber was then rinsed with 1 mL of HKEM buffer (50 mM KCl, 15 mM HEPES pH=7.5, 5 mM MgCl_2_, 1 mM EGTA).

Actin polymerization and contraction were induced by injection in the flow chamber of a solution containing 1μM actin monomers (12% labeled with Alexa 568), 3μM profilin, 25nM Arp2/3 complex, and 14nM of HMM-myosinVI (GFP labeled). This protein mixture was diluted in freshly prepared buffer containing 10mM Hepes (pH 7.5), 3mM ATP, 27mM DTT, 1mM EGTA, 50mM KCl, 5mM MgCl_2_, 3mg/ml glucose, 20μg/ml catalase, 100μg/ml glucose oxidase, and 0.25%w/v methylcellulose. The sample was then closed with VALAP to avoid evaporation.

### Patterning for heterogeneity experiments

Full, symmetrical and asymmetrical patterns were generated on a Nikon eclipse inverted microscope equipped with the Primo DMD (Alveole) (91). A flow chamber made with silanePEG 30k treated glass and cover glass and spaced with a 70μm double tape was filled with a photo initiator (PLPP). Patterns were designed in Inkscape and loaded into the μmanager’s Leonardo plugin (Alveole). The burning was done at 90mJ at 100% of a 5,6mW 365 nm wavelength laser. The flow chamber was then washed with 600μl of water to remove the PLPP and 200μl of SUV Buffer. 40μl of SUV solution was then added and incubated for 10 minutes at room temperature for an effective lipid coating. The excess of SUV was washed out by passing 600μl of SUV Buffer. The flow chamber was then passivated with 50μl of 1% BSA diluted in 1x HKEM. For the symmetrical and asymmetrical patterns, where a second round of patterning is required, 40μl of PLPP was injected. After a precise alignment, the burning was done at 70mJ at 80% of the laser. The PLPP was removed with 600μl of SUV buffer and equilibrated with 200μl of 1x HKEM. A mixture of 2 NPF (5nM Strep-WA, 1μM GST-PWA, 0,1% BSA in 1x HKEM) was injected and the coating was done by letting incubate for 15 minutes at room temperature. The excess of NPFs was washed out by passing 600μl of 1x HKEM and 30μl of the reaction mixture 0,5% BSA, 25nM Arp2/3, 3μM Human Profilin, 10nM GFP-Myosin VI, 1μM Actin 12% alexa 568 labeled in Fluorescence buffer (10mM Hepes (pH 7.5), 3mM ATP, 27mM DTT, 1mM EGTA, 50mM KCl, 5mM MgCl_2_, 3mg/ml glucose, 20μg/ml catalase, 100μg/ml glucose oxidase, and 0.25%w/v methylcellulose) was injected. The flow chamber was sealed with VALAP (Vaseline, Lanolin, Parafin 1:1:1) to avoid evaporation.

### Imaging

Time course of actin assembly and contraction on homogeneous patterns was acquired on a total internal reflection fluorescence (TIRF) microscope (Roper Scientific) equipped with an iLasPulsed system and an Evolve camera (EMCCD 512 × 512, pixel = 16 μm) using a 60× 1.49 objective lens. Microscope and devices were driven by MetaMorph software (Molecular Devices). For heterogeneous patterns, image acquisition was done on a Nikon Eclipse Ti2 inverted microscope equipped with a S Plan fluor ELWD 40x/0,60 objective and a Hamamatsu ORCA Flash 4.0 LT camera. Microscope and equipment were driven by MicroManager software.

### Image analysis

Image analysis was performed using Fiji (92), Rstudio and GraphPad Prism.

#### Analysis of contraction Area

Images were cropped according the size of the pattern and then binarized with Otsu method. Actin network area was determined with the “Analyse particles” function. On heterogeneous patterns, we determined and analyzed the actin area of each part of the pattern.

#### PIV Analysis

PIV analysis was performed with the “iterative PIV plugin” (93). The movie was first cropped around the pattern. We did the analysis on pairs of images taken every 3 frames in the different movies. Then, to analyze the PIV results, we separated the square pattern in four “quadrants” corresponding to the four corners of the square pattern. For each time of the PIV analysis, in each of these quadrants, we summed the vectors to obtain the value of the “PIV displacement” in each quadrant at each time point. Then we plotted this norm of vector sum for each quadrant as a function of time for networks contracting on glass or on lipids.

#### Myosin spots tracking

Myosin spots were detected and tracked with TrackMate plugin (94).

### Statistical analysis

We used Mann-Whitney tests (non-parametric test) performed in GraphPad Prism software.

### Computational modeling

We model the actomyosin network as a contractile viscoelastic medium represented by a lattice of actin and myosin nodes connected to each other by spring-like links, and to the substrate by adhesive dashpots. The network’s deformations are simulated numerically by solving force balance equations. Details of the model and numerical solutions are in the SI.

## Acknowledgements

We thank C. Copos for help with modeling. This work was supported by the European Research Council (Consolidator Grant 771599 (ICEBERG) to MT and Advanced Grant 741773 (AAA) to LB). This work was also supported by the MuLife imaging facility, which is funded by GRAL, a program from the Chemistry Biology Health Graduate School of University Grenoble Alpes (ANR-17-EURE-0003). MS and AM were supported by the National Science Foundation grant DMS 1953430 to AM. EMDLC was supported by the National Institutes of Health through award R35-GM136656.

