## Supplementary Information for "Friction patterns guide actin network contraction"

#### Supporting information

This file includes:

Supporting text (description of the model, pages 2-13)

Figures S1 to S5 (pages 14-19)

Legends for Movies S1 to S19 (pages 20-22)

Supporting References (page 23)

### Supporting Text

##### Mechanical model of contractile adhesive actomyosin network

_Qualitative model description_

We model the actomyosin network as a two-dimensional active viscoelastic material. It is well established that actin networks exhibit viscoelastic properties, with elastic behaviors predominantly seen at short timescales (seconds to tens of seconds) and viscous effects at long timescales (tens of seconds and longer) due to actin filament turnover and stochastic binding dynamics of crosslinks between filaments (1, 2). So, we model the system as elastic with relaxation (such that elastic stresses dissipate with time) and with active myosin contraction. The relaxation leads to viscous behavior on longer timescales. We also consider a nonlinear elasticity where the response to compression is disproportionately weaker than the response to stretch (see below). The deforming actomyosin network in the model experiences an effective viscous drag (friction) relative to the substrate.

The actomyosin mesh is coarse-grained as a triangular network of nodes connected by approximately equal length links. The links consist of a Maxwell element (elastic spring and viscous dashpot) and a contractile element in series (Figure 5A). The contractile elements yield a contractile tension determined by the myosin distribution. Each node is connected to the substrate by a viscous dashpot representing the friction between the network and substrate.

_Elastic Forces_

Consider a node $i$, linked to a neighboring node $j$. The link is modeled as an elastic spring that has rest length $\ell_{ij}^0$ and current length $\ell_{ij} = |\boldsymbol{x_i} - \boldsymbol{x_j}|$. The force magnitude exerted on node $i$ by the spring connecting node $i$ to node $j$, in series with a contractile element, is

$$F_{ij}^e = \gamma_{ij} + k_{ij}\left(\frac{\ell_{ij}}{\ell_{ij}^0} - 1\right) \qquad (1)$$

This force points from node $i$ to node $j$, along the unit vector $\boldsymbol{\tau_{ij}}$ connecting the nodes. The first term $\gamma_{ij}$ is the active contractile $(\gamma_{ij} > 0)$ force generated by myosin. Given a normalized myosin distribution at a given time $m(\boldsymbol{x})/m_0$, nodes are assigned 'myosin weights' $\gamma_i = \gamma_0 m(\boldsymbol{x})/m_0$ and the links between nodes are assigned the force equal to average of these weights: $\gamma_{ij} = (\gamma_i + \gamma_j)/2$. This is a simple way to capture the dependence of the contractile force from a myosin distribution on nodes.

The second term in Eq. (1) is a spring force, which depends only on the deviation of the current length $\ell_{ij}$ from the rest length $\ell_{ij}^0$. Following (3), the spring constant $k_{ij}$ has units of force. In the case of a linear elastic isotropic material, this link stiffness can be related to the material elastic modulus $\lambda_E$ (recall that in 2D, elastic moduli are reported with units $\sim pN/nm$) by equating the discrete strain energy with the continuous strain energy density in the limit of an infinitesimal deformation (3). For a regular mesh of equilateral triangles, such a computation yields a link spring constant

$$K_{ij} = \frac{8\lambda_E}{3\ell_{ij}^0}\left(\frac{dA_i + dA_j}{2}\right) \qquad (2)$$

Here, $dA_i$ is the area weight of node $i$ (the sum of one third of the area of each triangle with vertex $\boldsymbol{x_i}$). In this framework, the associated force density at node $i$ due to the spring between nodes $i$ and $j$ would be $\widetilde{F_{ij}^e} = F_{ij}^e / dA_i$.

We go beyond just Eq. (2) to model the network as nonlinearly elastic. Specifically, we assume that the spring constant is a function of the ratio $\ell_{ij}/\ell_{ij}^0$, as follows:

$$k_{ij} = \begin{cases} K_{ij}, & \frac{\ell_{ij}}{\ell_{ij}^0} - 1 \geq 0 \\ r_k K_{ij}, & \frac{\ell_{ij}}{\ell_{ij}^0} - 1 < 0 \end{cases} \tag{3}$$

Thus, the spring responds to stretching ($\ell_{ij}/\ell_{ij}^0 > 1$) and compression ($\ell_{ij}/\ell_{ij}^0 < 1$) differently. Here $K_{ij}$ is initialized as in Eq. (2). Parameter $r_k$ is strictly less than 1, so the network can be easily contracted but resists stretching. This nonlinear feature is inspired by modeling the actin mesh as a 'cable' network (4): individual actin filaments and bundles of filaments are not stretchable but can easily bend or buckle.

*Force Balance*

The internal network forces (active myosin stress and passive elastic forces) are balanced by the friction with the substrate, which is characterized by a drag coefficient $\zeta$ which may depend on position. The governing force balance equation for node $i$ is then:

$$\sum_{\text{neighbors } j} \left( \gamma_{ij} + k_{ij} \left( \frac{\ell_{ij}}{\ell_{ij}^0} - 1 \right) \right) \boldsymbol{\tau_{ij}} = \zeta \, dA_i \frac{d\boldsymbol{x_i}}{dt} \tag{4}$$

Note that the drag on a node $i$ depends also on its area weight $dA_i$ – essentially, we assume the effective drag on a node to be proportional to its actin density. Since the bulk network is initialized to be uniform, the initial area weight $dA_i$ is a proxy for actin density. So thus, a node with a higher actin density or approximating a larger portion of the network (both yielding more area weight $dA_i$) should experience more effective drag.

*Network relaxation*

Thus far, Eq. (1) – (4) describe a nonlinear elastic network. We model the relaxation of the elastic stresses, represented by the dashpot element of links in main text Fig. 5a, by the relaxation of the rest length of elastic springs $\ell_{ij}^0$ to their current length with rate $r$:

$$\frac{d\ell_{ij}^0}{dt} = r(\ell_{ij} - \ell_{ij}^0) \tag{5}$$

For example, if the current length is fixed to be larger than the rest length $\ell_{ij} > \ell_{ij}^0$, then over time $\ell_{ij}^0$ will grow until it reaches $\ell_{ij}$. Similarly, if the current length is shorter than the rest length $\ell_{ij} < \ell_{ij}^0$, then the rest length will shrink until it reaches $\ell_{ij}$.

Since the link spring constants $k_{ij}$ are initialized using Eq. (2) and (3), there is an inherent dependence of the spring constant on the rest lengths local to a node. Presuming that throughout time links are approximately equal locally, this dependence is approximated as $k_{ij} \propto \ell_{ij}^0$. So, as the rest length $\ell_{ij}^0$

changes with time, the value $k_{ij}/\ell_{ij}^0$ is kept constant for each link. This ensures that the global elastic properties of the network are approximately constant over time, despite relaxation effects.

*Myosin contractile forces*

Most of the myosin concentration is highly localized into several aggregates (spots). One or several myosin foci appear at locations that we glean from the experiments. Each myosin foci generates radially symmetric stress that rapidly decreases with distance from the center of the foci (in other words, the stress is relatively local). We consider the following simple description for the myosin-generated stress - a myosin spot which initially forms centered at node $i_M$. This spot center is advected with the network, meaning it is always attached to node $i_M$ throughout the deformation process. At any given time, links in a region of characteristic radius approximately equal to $2\sigma_M$ experience a nonzero tension $\gamma_{ij}$. As previously mentioned, each node $j$ is assigned a 'myosin weight' $\gamma_j$. For a single myosin spot, this weight $\gamma_j$ is:

$$\gamma_0 \exp\left(-\left(\frac{x_j - x_{i_M}}{\sigma_M}\right)^2 - \left(\frac{y_j - y_{i_M}}{\sigma_M}\right)^2\right) \qquad (6)$$

The links between nodes are then assigned the force equal to average of these weights: $\gamma_{ij} = (\gamma_i + \gamma_j)/2$. Nodes that get too close to the node $i_M$ carrying the myosin spot are absorbed, with their area weight $dA_j$ added to the myosin spot, effectively increasing the drag between this node and substrate. Thus, over time, all the area weight ($\sim$actin density) should converge to the myosin spot node (which itself will move with time).

For simplicity, in the simulations, all myosin spots are identical and scale as $\gamma_0$ divided by the number of the initial spots (this way, the total myosin strength is approximately constant in cases considered). However, see additional notes below about the myosin stresses.

*Comments on the network's rheology*

The internal material properties of the network are not known in detail, and in principle, on the relevant time scale longer than minutes, the network could be purely viscous. However, when we simulated the network as a compressible viscous fluid, the resultant velocity fields for a given myosin distribution were different from the observations. Though the boundary conditions and inherent boundary shape would result in flow toward the center in homogeneous patterns, the relative magnitudes of local network flow (and thus resultant deformation) close to the myosin spot and far from it were different from the observations. Most importantly, flow between myosin spots was not quiescent as observed in experiments (see Figure. 3). On the other hand, a purely elastic network, of course, only deforms partially and never compacts to the center. So, a viscoelastic description is necessary.

Lastly, having a network more resistant to stretching works best because network compaction close to myosin spots is more effective than if resistance to compression is equally high. More importantly, we know resistance to stretch is required for capturing the observed lack of flow between myosin spots and having a higher resistance to stretch than compression best captures the ratio of flow magnitudes between spots versus close to spots being much smaller than 1. All these reasons and trial simulations with other possible rheologies provided rationale for the model choice.

**Parameters**

Non-dimensional model parameters are listed in this table:

| **Parameter** | Non-dimensional value |
|---|---|
| *Elastic modulus* $\lambda_E$ | 50 |
| *Discretization length* $h_x$ *(approximate starting link length*) | $1/16 \ = 0.0625$ |
| *Stretch spring constant* $$k_{ij} = \frac{8\lambda_E}{3\ell_{ij}^0}\left(\frac{dA_i + dA_j}{2}\right)$$ | Initialized as $\sim$ 8 for internal nodes, between $\sim$ 3 and $\sim$ 5 for border nodes, when using the above $\lambda_E$ and $h_x$ |
| *Ratio of compression to stretch spring constant* $r_k$ | 0.15 |
| *Relaxation time* $1/r$ | 1/2 |
| *Maximum contractile force* $\gamma_0$ | 2.0 |
| *Contraction range* (gaussian width $\sigma_m$) | 0.05 |
| *Lipid drag coefficient* $\zeta_1$ | 300 |
| *Glass drag coefficient* $\zeta_2$ | 850 |
| *Timestep* $dt$ | 0.0001 |

These choices are justified as follows. The non-dimensional length scale is such that one length unit corresponded to the width of the pattern (square width, 60 $\mu m$, or disk diameter, 70 $\mu m$, which is also used for the rectangle case). The initial rest length of the springs is set to be roughly equal to the initial discretization length which is 1/16th of the non-dimensional length unit. To operate with forces on the order of unity, it is convenient to have the compression spring constant on the order of unity. We found that this is the case if we choose the non-dimensional value for the elastic modulus equal to 50. The ratio $k_{ij}/\ell_{ij}^0$ was kept constant throughout the simulations. The stretch spring constant must be at least a few fold greater than the compression spring constant for the global stress propagation in the network, but not more than an order of magnitude greater to avoid numerical instabilities associated with occasional local remeshing which produces relaxed links. We found that the compression / stretch spring constant ratio 0.15 worked well in combination with compression spring constants initialized to be at least $\sim 0.5$. We chose the maximal contractile force to be on the order of unity; then, at the observed strains of the network and chosen relaxation rate, respective internal elastic forces are approximately a magnitude smaller than the local myosin contractile forces, and so the local contractile behavior is predominantly the balance between the myosin active force and the friction between the network and substrate. To keep the myosin contraction local and highly concentrated in spots, we chose the contraction range to be on the order of the average node-to-node distance. We chose the order of magnitude of the non-dimensional viscous drag (per small area around the node) coefficients for this friction in the range from 100 to 1000;

given the chosen discretization size $h_x$, this means the product of drag and area weight $\zeta dA_i$ on a node is usually on the $O(0.1 - 1.0)$ (so, of a comparable magnitude to forces considered). The drag coefficient indirectly determines the time unit (because the force and length units are already chosen, and viscous drag's dimension contains units of force, length and time), so choosing a larger non-dimensional viscous drag, given the choice of timestep $dt$, allows for the contraction to proceed slowly enough to avoid numerical problems but not so slowly that computation runtime is exceedingly high. The time step was necessitated by considerations of numerical stability. With these parameters, the network compaction (on lipid) was generally completed in ~ 10 non-dimensional time units (approximately 13 non-dimensional time units on the square pattern), which we took to correspond to 23 min of biological time to match the measurements. The links' characteristic relaxation time was chosen to be on the order of one non-dimensional time unit to be much shorter than the duration of the network compaction, in order for the long-term network behavior to be more viscous than elastic, but not smaller than one non-dimensional time unit in order for the elastic stresses not to become too weak too fast, which would make the contraction very local. Additionally, the relaxation time cannot be so short that stretched links are instantly relaxed, which would mean the disproportionate stretch to compression response would not be evident. Lastly, the ratio of the glass to lipid viscous drags was determined by fitting the ratio of the resulting rates of area decrease for various choices of viscous drag coefficients, such that the relative rates of area decrease match experimental quantitative estimates.

**Numerical implementation**

*Initialization*

Given a chosen domain shape (square, disk, rectangle), the domain is coarse-grained using the distmesh package (5) which utilizes an algorithm which employs Delaunay triangulation to generate a network of points with links of $\approx h_x$ equal length. The initial links' lengths are assigned as the starting rest lengths, such that initially the network has no passive internal stresses, and if no external or active forces are applied then the material does not deform.

*Time-stepping*

At each timestep, links are assigned their active contractile tensions $\gamma_{ij}$ based on the sum of myosin spot contributions given by Eq. (6). The elastic forces are calculated, and node positions $\boldsymbol{x_i(t)}$ are updated by taking a Forward Euler step of Eq. (4). Simultaneously, the link rest lengths $\ell_{ij}^0$ are updated also with a Forward Euler step of Eq. (5), and the spring stiffnesses are accordingly adjusted as previously described.

*Myosin spot node absorption*

We analyzed the quantity of myosin in foci compared to the total intensity of myosin (not shown). During the contraction phase, ratio of myosin in foci to total quantity of myosin is high and remains close to one over time. This is why in simulations we focused on the myosin localized within foci.

The effect of a myosin spot is primarily localized within a $2\sigma_M$ distance of node $i_M$. This parameter $\sigma_M$ is chosen to be comparable to the initial node-node distance $h_x$. When a node $j$ that is a first neighbor of the myosin spot and within a $h_x/6$ distance from the spot, node $j$ is "absorbed" by node $i_M$ such that $dA_{i_M} \to dA_{i_M} + dA_j$. This absorption distance is chosen such that there is little difference in the calculation of contractile forces $\gamma_{ij}$ locally with or without node $j$. The myosin spot and other remaining first neighbors of node $j$ become connected with a local Delaunay triangulation, assigning the new links rest lengths $\ell_{ij}^0$ which are relaxed ($= \ell_{ij}$) and spring constants $k_{ij}$ approximately equal to that of the lost edges. This choice of $\ell_{ij}^0$ and $k_{ij}$ for the new edges ensures any transient responses due to remeshing are minimized – assigning the rest lengths and spring constants differently from this would be relatively inconsequential, since the contractile forces $\gamma_{ij}$ on links close to the myosin spot is the dominant internal force for the nodes.

*Remeshing and extra numerical conditions*

It is necessary to add additional implementation measures to prevent unphysical behaviors (for example, with strong enough non-uniform compressive loads the network could overlap over itself). Since there is long-time viscous behavior, it is appropriate to occasionally perform local remeshing of the network to delete and/or add new links to prevent these behaviors. We implement two kinds of local remeshing: remeshing of low-quality triangles (as determined by distmesh function simpqual) and remeshing of cusped boundary points.

If the quality of a triangle, which ranges from $0$ to $1$, is $< 0.05$ then this triangle and all its nodes' first neighbors are remeshed according to Delaunay triangulation rules. No links which overlap the rest of the triangulation are added. These new edges have rest lengths $\ell_{ij}^0$ which are relaxed with spring constants $k_{ij}$ equal to the average lost edge spring constant times the ratio of the new rest length to average old edge rest lengths. This ensures that the network does weaken due to local remeshing, while also minimizing any transient response due to remeshing. The low-quality triangle remeshing occurs until at most 40% of the network becomes localized at the myosin spots, which is generally when most of the network has contracted to a small region and nearly all triangles become low quality.

At initialization, the boundary nodes of the network are known. Considering these nodes as forming a polygon, the exterior angles are tracked with time. If an exterior angle drops below $100°$, as typically occurs when a piece of the domain boundary approaches a myosin spot, the associated exterior node is removed and its area weight is redistributed to its neighbors equally. The node's first neighbors are reconnected according to Delaunay triangulation rules, proceeding as described above for low quality triangles. The boundary remeshing occurs until at most 80% of the network becomes localized at the myosin spots, after which very few boundary nodes remain and the network is highly compacted so remeshing is unnecessary.

In the rare instances that a node disconnects from the bulk network due to these local remeshing rules, it is removed from the system and its area weight is redistributed to its closest neighbors. Generally, this occurs rarely and in regions when nodes are densely packed, so any choice of node weight distribution does not significantly affect even the quantitative results.

Finally, if a node pair has link distance $\ell_{ij} < h_x/11$, we assume the spring constant $k_{ij}$ becomes infinite so that the link cannot compress any further. We found that a condition of this form is necessary to avoid unphysical behaviors close to the myosin spots, where contraction is very high. Effectively, this condition ensures that such node pairs move as a unit together until one or both are absorbed by a myosin spot.

**Location of the network convergence point**

On a uniform pattern, the network compacts to the centroid.

At every node $i$, the internal forces (active contraction, elastic responses, etc.) are balanced with the external friction force:

$$\boldsymbol{F}_{\boldsymbol{i}}^{\boldsymbol{int}}(\boldsymbol{x}_i) = \zeta(\boldsymbol{x}_i)\frac{d\boldsymbol{x}_i}{dt}dA_i \quad (7)$$

where $\boldsymbol{x}_i(t)$ is the position of the node $i$ over time and $dA_i$ is the area weight, a proxy for the initial actin density carried by the node. Note that we presume the actin density to be fixed and advected on the nodes. For simplicity, let's assume that all nodes carry the same initial actin mass, taking $dA_i = 1$ in Eq. (7). The internal forces over all nodes must sum to zero according to Newton's third law (as an internal force on node $i$ from node $j$ is equal and opposite to the internal force on node $j$ from node $i$). This means that

$$\sum_i \boldsymbol{F}_{\boldsymbol{i}}^{\boldsymbol{int}}(\boldsymbol{x}_i) = \boldsymbol{0}$$

Necessarily, by applying this equality to Eq. (7) we then have:

$$\sum_i \zeta(\boldsymbol{x}_i)\frac{d\boldsymbol{x}_i}{dt} = \boldsymbol{0} \quad (8)$$

First, consider Eq. (8) in the case of a homogeneous pattern where $\zeta(\boldsymbol{x}) = \zeta$, a constant. We can swap the time derivative with the summation,

$$\frac{d}{dt}\sum_i \zeta\, \boldsymbol{x}_i = \boldsymbol{0}$$

then time integrating gives us

$$\sum_i \zeta\, \boldsymbol{x}_i = \boldsymbol{const}.$$

Equivalently

$$\sum_i \zeta\, \boldsymbol{x}_i / \sum_i \zeta = \boldsymbol{const}. \quad (9)$$

Notice that the left-hand-side of Eq. (9) is a center-of-mass calculation, except mass density is replaced by viscous drag. More generally when drag may depend on node position, we refer to summations of the form

$$\sum_i \zeta(\boldsymbol{x}_i)\, \boldsymbol{x}_i / \sum_i \zeta(\boldsymbol{x}_i) \quad (10)$$

as the instantaneous center-of-drag of the system.

Eq. (9) then states that at any stage of the contraction, the center-of-drag of the network does not shift. Necessarily, the compaction point at which all $\boldsymbol{x}_i$ node positions converge is exactly the center-of-drag at the final time, which by Eq. (9) equals the center-of-drag at the onset. Though the node positions $\boldsymbol{x}_i = \boldsymbol{x}_i(t)$ are time dependent and may vary based on myosin distribution, we can easily compute the center-of-drag at the contraction onset. Since we are dealing with the especially straightforward case of a homogeneous pattern, this center-of-drag coincides with the centroid: the geometric center of the chosen domain. This is exactly the convergence point we observe in the homogeneous square, disk, and rectangle cases.

Now, suppose that the pattern is not homogeneous, i.e. $\zeta(\boldsymbol{x})$ depends on the position $\boldsymbol{x}_i$ of a node. It is easiest to discuss this in the context of viewing the network as a continuum, continuing in Lagrangian coordinates – meaning $\boldsymbol{x} = \boldsymbol{x}(\boldsymbol{x_0}, t)$ where $\boldsymbol{x_0}$ is the initial position in the reference configuration. The equivalents to Eq. (7) and (8) are:

$$\boldsymbol{F^{int}}(\boldsymbol{x}) = \zeta(\boldsymbol{x})\frac{dx}{dt} \qquad (7')$$

$$\int \zeta(\boldsymbol{x})\frac{dx}{dt}\, dA_0 = 0 \qquad (8')$$

Here $\boldsymbol{F^{int}}(\boldsymbol{x})$ is the force density. In the context of patterns which are not homogeneous, we seek to understand how the center of drag

$$\boldsymbol{c}_D = \int \zeta(\boldsymbol{x})\boldsymbol{x}\, dA_0 \,/\! \int \zeta(\boldsymbol{x})\, dA_0 \qquad (11)$$

changes with time. Note that Eq. (11) is the continuous analogue of Eq. (10). The final position of the center of drag $\boldsymbol{c}_D$ must necessarily coincide with the convergence point of the nodes. Taking the time derivative of Eq. (11), we have

$$\frac{d}{dt}(\boldsymbol{c}_D) = \frac{d}{dt}\left(\int \zeta(\boldsymbol{x})\boldsymbol{x}\, dA_0\right)\frac{1}{\int \zeta(\boldsymbol{x})dA_0} + \int \zeta(\boldsymbol{x})\boldsymbol{x}\, dA_0\left(-\frac{\frac{d}{dt}\left(\int \zeta(\boldsymbol{x})dA_0\right)}{\left(\int \zeta(\boldsymbol{x})dA_0\right)^2}\right)$$

$$= \frac{1}{\int \zeta(x)dA_0}\left(\int \zeta(\boldsymbol{x})\frac{dx}{dt}\, dA_0 + \int \nabla\zeta\cdot\frac{dx}{dt}\,\boldsymbol{x}\, dA_0 - \frac{\int \zeta(x)x\, dA_0}{\int \zeta(x)dA_0}\int \nabla\zeta\cdot\frac{dx}{dt}\, dA_0\right) \qquad (12)$$

The first term in Eq. (12) is identically zero by Eq. (8’), so we have that the center of drag changes with time as:

$$\frac{d}{dt}(\boldsymbol{c}_D) = \left(\int \nabla\zeta\cdot\frac{dx}{dt}\,\boldsymbol{x}\, dA_0 - \boldsymbol{c}_D\int \nabla\zeta\cdot\frac{dx}{dt}\, dA_0\right)\Big/\!\int \zeta(\boldsymbol{x})dA_0 \quad (13)$$

In particular, notice the relevance of the inner product of the change in drag with the velocity; this term $\nabla\zeta\cdot\frac{dx}{dt}$ is nonzero when there is flow parallel to the direction of change in drag.

To gain further intuition as to the meaning of Eq (13), let us consider the case of a heterogeneous rectangular domain, where

$$\zeta(x,y) = \begin{cases}\zeta_1, & x < 0\\ \zeta_2, & x \geq 0\end{cases} = \Delta\zeta\, H(x) + \zeta_1$$

Let us choose $\Delta\zeta = \zeta_2 - \zeta_1 > 0$ (as is consistent with a lipid-drag left-right divide). Note that $H(x)$ is the Heaviside step function. Then, in this pattern case, the gradient of drag is simply

$$\nabla\zeta = <\Delta\zeta\, \delta(x), 0>$$

Then, letting $\frac{dx}{dt} = <u, v>$, Eq. (13) becomes

$$\frac{d}{dt}(\boldsymbol{c}_D) = \frac{1}{\int \zeta(x)dA_0}\left(\int \Delta\zeta\,\delta(\mathrm{x})\, u\,\boldsymbol{x}\, dA_0 - \boldsymbol{c}_D\int \Delta\zeta\,\delta(\mathrm{x})\, u\, dA_0\right) \quad (14)$$

Because of the delta function in $x$, integrals are only nonzero when considering reference positions $\boldsymbol{x_0}$ such that $x = 0$ (i.e. we only integrate over particles which currently at time $t$ live on the lipid-drag divide). Let us call the set of reference positions who's current $x$ position is $x = 0$ the set $S(t) = \{\boldsymbol{x_0} \text{ such that } x(\boldsymbol{x_0}) = 0\}$. Looking more closely at the time evolution of $\boldsymbol{c_D} \cdot x$, since that is the direction with asymmetry, we have

$$\frac{d}{dt}(\boldsymbol{c_D} \cdot \hat{\boldsymbol{x}}) = \frac{-\boldsymbol{c_D} \cdot \hat{\boldsymbol{x}}}{\int \zeta(x) dA_0} \int_{S(t)} \Delta\zeta \, u \, dA_0 \quad (15).$$

Now, remember $\Delta\zeta > 0$ and we can assume that the horizontal flow $u$ along the boundary $x = 0$ is nearly always to the right, i.e. $u > 0$. The total drag at any moment in time, $\int \zeta(\boldsymbol{x}) dA_0$, is also strictly positive. Finally, the $x$ coordinate of the center of drag $\boldsymbol{c_D} \cdot \hat{\boldsymbol{x}}$ is expected to be preferentially in the $x > 0$ regime, since the $x > 0$ side has a higher drag coefficient than the $x < 0$ side of the regime. All this put together suggests that, in fact,

$$\frac{d}{dt}(\boldsymbol{c_D} \cdot \hat{\boldsymbol{x}}) < 0 \quad (16)$$

for the majority of time $t$ during contraction. Eq. (16) means we expect the $x$ coordinate of the center of drag $\boldsymbol{c_D}$ to shift away from its $t = 0$ value in the direction of the lipid-glass divide where the drag changes.

Given the simple case of a heterogeneous domain, we have gained valuable intuition regarding Eq. (13), which more generally describes how the center of drag changes with time. Eq. (13) tells us that there are two primary factors: one, the "drag flux" associated with the flow in the direction of changes in drag coefficient spatially and two, the location of these spatial changes in the drag coefficient (as is suggested by first term $\int \nabla\zeta \cdot \frac{dx}{dt} \, \boldsymbol{x} \, dA_0$).
More generally, in the heterogeneous case, the movement of nodes into regimes of higher drag penalizes the center of drag preferentially in the direction of this change.

*Estimate of the convergence point coordinates on the heterogeneous pattern*

We have just shown that the cross-over of the network into regions of higher drag penalizes the center of drag towards the lipid-drag divide, but what is less clear is the extent to which the center of drag moves from its starting location at time $t = 0$.

Let us first calculate the initial center of drag $\boldsymbol{c_D}(t = 0)$, as in Eq. (11), in all considered experimental cases. This initial position should be an upper bound of sorts on the final $\boldsymbol{c_D}$ after compaction. We consider the square $[-0.5,0.5]^2$, rectangle $[-0.75,0.75] \times [-0.25,0.25]$, and disk centered at 0 with radius 0.5 (here, 1 is the non-dimensional length unit). Let the lipid drag coefficient be $\zeta_1$ and the glass drag coefficient be $\zeta_2$. Then,

$$\boldsymbol{c_D}(t = 0) = < 0.25 - \frac{1}{2}\left(1 + \frac{\zeta_2}{\zeta_1}\right)^{-1}, 0 > \quad \text{[heterogenous square]}$$

$$\boldsymbol{c_D}(t = 0) = < \frac{1}{3}\left(\frac{1+2\,\zeta_2/\zeta_1}{1+\zeta_2/\zeta_1}\right) - 0.5, \frac{1}{3}\left(\frac{1+2\,\zeta_2/\zeta_1}{1+\zeta_2/\zeta_1}\right) - 0.5 > \quad \text{[diagonal square]}$$

$$\boldsymbol{c_D}(t = 0) = < 0.375 - 0.75\left(1 + \frac{\zeta_2}{\zeta_1}\right)^{-1}, 0 > \quad \text{[heterogenous rectangle]}$$

$\boldsymbol{c}_D(t=0) =< \frac{1}{2}\left(\frac{1+2\,\zeta_2/\zeta_1}{1+\zeta_2/\zeta_1}\right) - 0.75, \frac{1}{6}\left(\frac{1+2\,\zeta_2/\zeta_1}{1+\zeta_2/\zeta_1}\right) - 0.25 >$ [diagonal rectangle]

$\boldsymbol{c}_D(t=0) =< \frac{2}{3\pi}\left(\frac{\zeta_2/\zeta_1-1}{\zeta_2/\zeta_1+1}\right), 0 >$ [heterogenous disk]

In all these cases, for times $t > 0$ we expect $\boldsymbol{c}_D$ to shift in the direction of the lipid-glass divide. Without additional factors in the dynamics, this puts a hard limit on the depth of the contraction point in the glass regime.

Since generally lipid surfaces tend to contract before drag surfaces, let us consider the special case where the lipid portion of the domain has entirely contracted to the lipid-glass boundary. There are experimental observations (Figure 4) that suggest this special case to be almost true since the lipid and glass surfaces have different lag times before contraction. Note that if the drag part of the network is fixed (effectively with a very large drag coefficient) while the lipid part is contracting, up until the glass part starts contracting the center of drag $\boldsymbol{c}_D$ will be farther into the glass portion of the domain than as calculated above. So even if $\zeta_2/\zeta_1$ is close to 1, the time delay in contraction inherently shifts the drag coefficient away from the centroid.

For this special case, let us presume that the lipid part of the network contracts to the drag boundary and that it is distributed along this boundary as if contractile flow was net perpendicular to the boundary. In the square and glass cases, this would yield a distribution of the compacted lipid portion which is symmetric around the center of the domain. Once the lipid portion of the network has completely contracted to the boundary and if we presume the glass portion of the network is uniform, then we can apply the center of drag argument for homogeneous domains, as in Eq. (9), as all of the network experiences the same drag coefficient. Note that, presuming the lipid contracts fully to the boundary first, then *regardless of the choice of drag coefficients* the convergence point of the network must be

$\boldsymbol{c}_D^*(t_{final}) =< 0.125, 0.0 >$ [heterogenous square]

$\boldsymbol{c}_D^*(t_{final}) =< 7/12, 7/12 > \approx < 0.083, 0.083 >$ [diagonal square]

$\boldsymbol{c}_D^*(t_{final}) =< 0.1875, 0.0 >$ [heterogenous rectangle]

$\boldsymbol{c}_D^*(t_{final}) = < 0.025, 0.075 >$ [diagonal rectangle]

$\boldsymbol{c}_D^*(t_{final}) =< 1/3\pi, 0 > \approx < 0.10, 0.0 >$ [heterogenous disk]

Note that excluding the rectangular diagonal case[1], $\boldsymbol{c}_D^*(t_{final})$and $\boldsymbol{c}_D(t=0)$ are identical for a glass/lipid ratio of $\zeta_2/\zeta_1 = 3$. Below this ratio, $\boldsymbol{c}_D(t=0) < \boldsymbol{c}_D^*(t_{final})$ and above this ratio $\boldsymbol{c}_D(t=0) > \boldsymbol{c}_D^*(t_{final})$. Essentially, this tells us that if the glass/lipid drag ratio is $\zeta_2/\zeta_1 < 3$ then a time delay in contraction pushes the final contraction position closer to $\boldsymbol{c}_D^*(t_{final})$, beyond $\boldsymbol{c}_D(t=0)$.

---

[1] In the diagonal rectangle case, we assumed the flow was perpendicular to the drag boundary. This preferentially places the lipid portion of the network to be primarily in the $x < 0$ part of the domain, while the glass portion is primarily in the $x > 0$ part of the domain due to the aspect ratio. This affects the final position be close to $x = 0$ but this is not necessarily what we would expect the effect of flow through the boundary to be on $\boldsymbol{c}_D(t > 0)$.

If the glass/lipid drag ratio is $\zeta_2/\zeta_1 > 3$, $\boldsymbol{c}_D^*(t_{final})$ should still be a rough estimate of the final contraction position, since the glass portion of the network is mostly stationary as the lipid part contracts so we can reasonably expect the lipid portion of the network to contract to the boundary before significant deformation of the glass part has occurred.

In sum, with our understanding of how the center of drag changes over time (Eq. (12)) and the estimated final positions $\boldsymbol{c}_D^*(t_{final})$ for the special case associated with a time delay, generally we expect that the final position be close to $\boldsymbol{c}_D^*(t_{final})$, except in the diagonal rectangular case where the predicted final position is more likely to be point along the line from the drag boundary to the initial center of drag $\boldsymbol{c}_D(t = 0)$.

*Why theory and simulation underestimate the distance between the convergence point and centroid on the heterogeneous pattern*

If the domain is heterogenous (lipid/glass), the numerical simulation predictions for the convergence points are slightly lower than the final analytical estimates of the previous section. This simulation underestimation is partly because the simulations rarely reach the extreme case of the lipid side fully contracting to the boundary (and there is always some minimal error associated with discretization and implementation simplifications). But even these analytical estimates of the distance between the convergence point and centroid of the pattern are slightly lower than the *average* experimentally measured distance (the lowest measured distances compare nicely to the computed ones). There are likely two reasons for the underestimation by the model. First, in the model we neglected network disassembly, but some disassembly likely does take place. Then, it is likely that the part of the network that is initially on lipid will start disassembling earlier because it started contracting earlier. This would give a lower 'weight' to the drag of the network on the lipid part, shifting the center of the drag deeper into the glass part. Second, the retraction fibers, clearly visible in the final stages of the network compaction, could be effectively pulled into the final large myosin aggregate at the focal point of these fibers' aster. The aggregate's position is then biased by the effective friction between these fibers and substrate. Under some simple assumptions, this bias is toward the center of the glass part of the pattern.

*Centering with acceleration*

Numerical simulations predict velocity of the myosin spots directed toward the centroid of the homogeneous pattern, and roughly toward the center-of-drag of the heterogeneous pattern, with speed gradually slowing down near the convergence point. The direction is predicted correctly, but the experimentally observed speed increases near the convergence point. The simple explanation of this phenomenon is: First, we observe that the total myosin amount in the myosin spots increases substantially during the contraction. From the model, the spots' speeds accelerate with the increase of the myosin strength, which is proportional to the myosin density. Second, the network likely undergoes partial disassembly during the contraction. Because of that, the friction force is likely to gradually decrease, also serving to accelerate the contraction.

*Square pattern contracts faster than disc pattern*

This observation can be explained by the model providing the following two assumptions are true: the average amount of myosin per spot does not depend on the number of spots (and so the total amount of myosin in all spots increases with the spot number), and the average number of spots per pattern is greater on a square than on a disc. The second of these assumptions is true, based on the observations. The reason for that is likely that the initial myosin spots preferentially appear in the corners of the pattern because initial myosin condensation is accelerated in the corner: distributed contraction is faster near the outwardly curving boundary, as previously published simulations predict (6). We also saw in our simulations that at the initial stage of contraction, each spot generates a roughly constant rate of the area decrease, and as far as the spots are far from each other, the effect is additive. Then, if the first assumption is correct, the net rate of the area reduction scales with the number of spots, explaining why the rate of the area decrease on the square pattern is greater than that on the disc pattern.

*Why centering error is greater on glass than on lipid*

We observed that on the glass square pattern, the initial distribution of the myosin spots is most often asymmetric (for example, there is one spot in one of the corners, or two spots in the adjacent corners). In those cases, due to the subtle factors not included into the model, like contraction-dependent partial network disassembly and/or some dependence of the effective friction on the history of drag across local patches of the surface, these factors would translate into asymmetries of the contractions violating the theorem of the compaction to the centroid. In other words, the more asymmetric the initial myosin spot distribution is, the greater the shift from the center of the pattern is for the convergence point, which is the case on the glass patterns. On the lipid square patterns, the initial myosin spots are distributed more symmetrically, like four spots in all four corners. In this case, the contraction must be symmetric, converging exactly to the center.

#### Supporting Figures

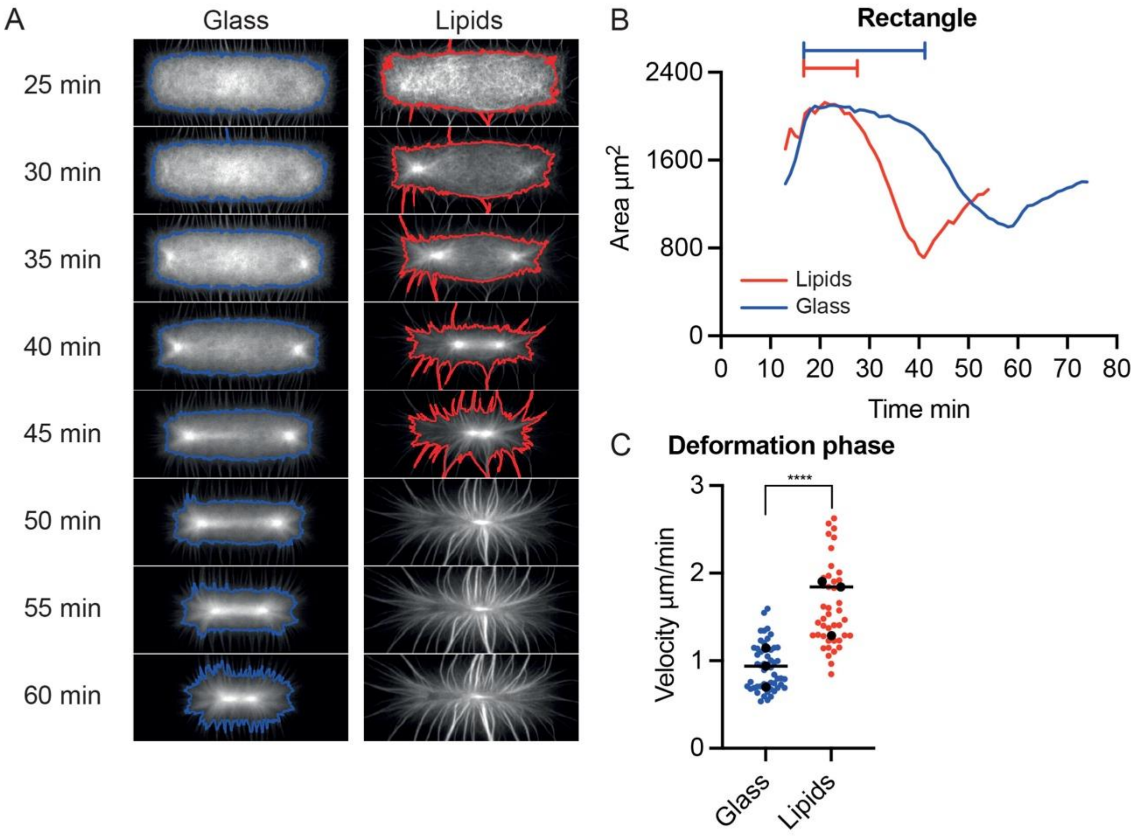


**Supporting Figure 1. Contraction of rectangle-shaped actin network on glass or lipid micropattern.**

**A.** Time-lapse imaging of actin network contraction on a glass or lipid rectangle (L=104µm, l=34µm) micropattern. Line (Blue for glass, Red for lipids) correspond to the contours of the actin network (see methods). Biochemical conditions: On glass micropattern, WA = 1 µM; on lipid micropattern: WA = 1 nM. Actin 1µM, Human Profilin 3µM, Arp 25nM, Myosin VI 14nM. **B.** Measured actin area as a function of time for the lipid (red) or glass (red) conditions. **C.** Velocity of the phase contraction phase for the lipid or glass conditions on rectangle micropattern. Data are represented with a superplot. Rectangle Glass n=52-N=3-*median* =0.9089, Rectangle Lipids n=68-N=4-*median* =1.474. Mann-Whitney Statistics : Rectangle Glass/Rectangle Lipids *p* value≤0.0001 ****.

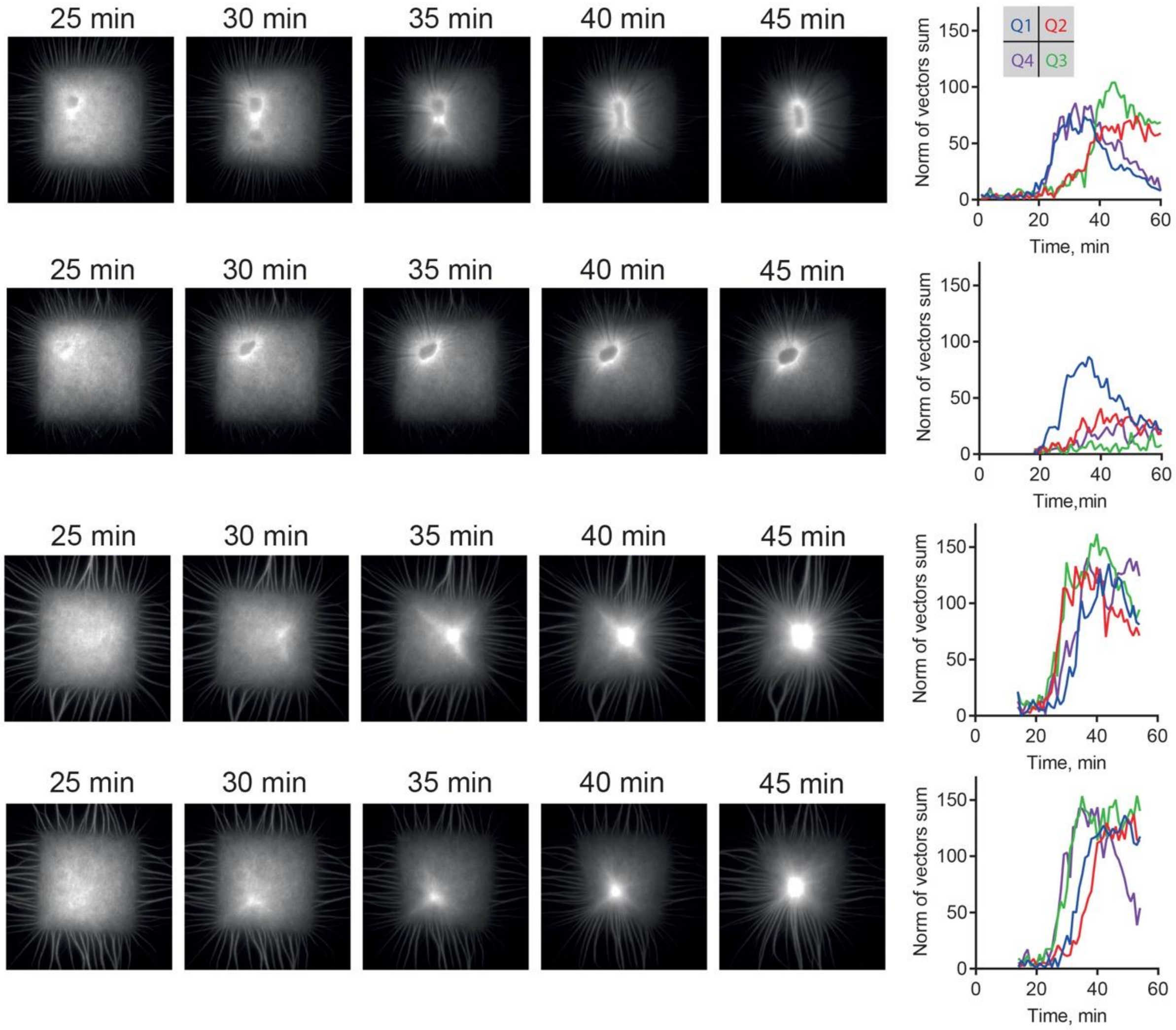


**Supporting Figure 2. Actin flux is heterogeneous and asymmetric on glass micropatterns.** Four examples of actin networks grown on glass substrate with the resultant of vector sum for each quadrant defined on the pattern as a function of time (obtained with PIV analysis).

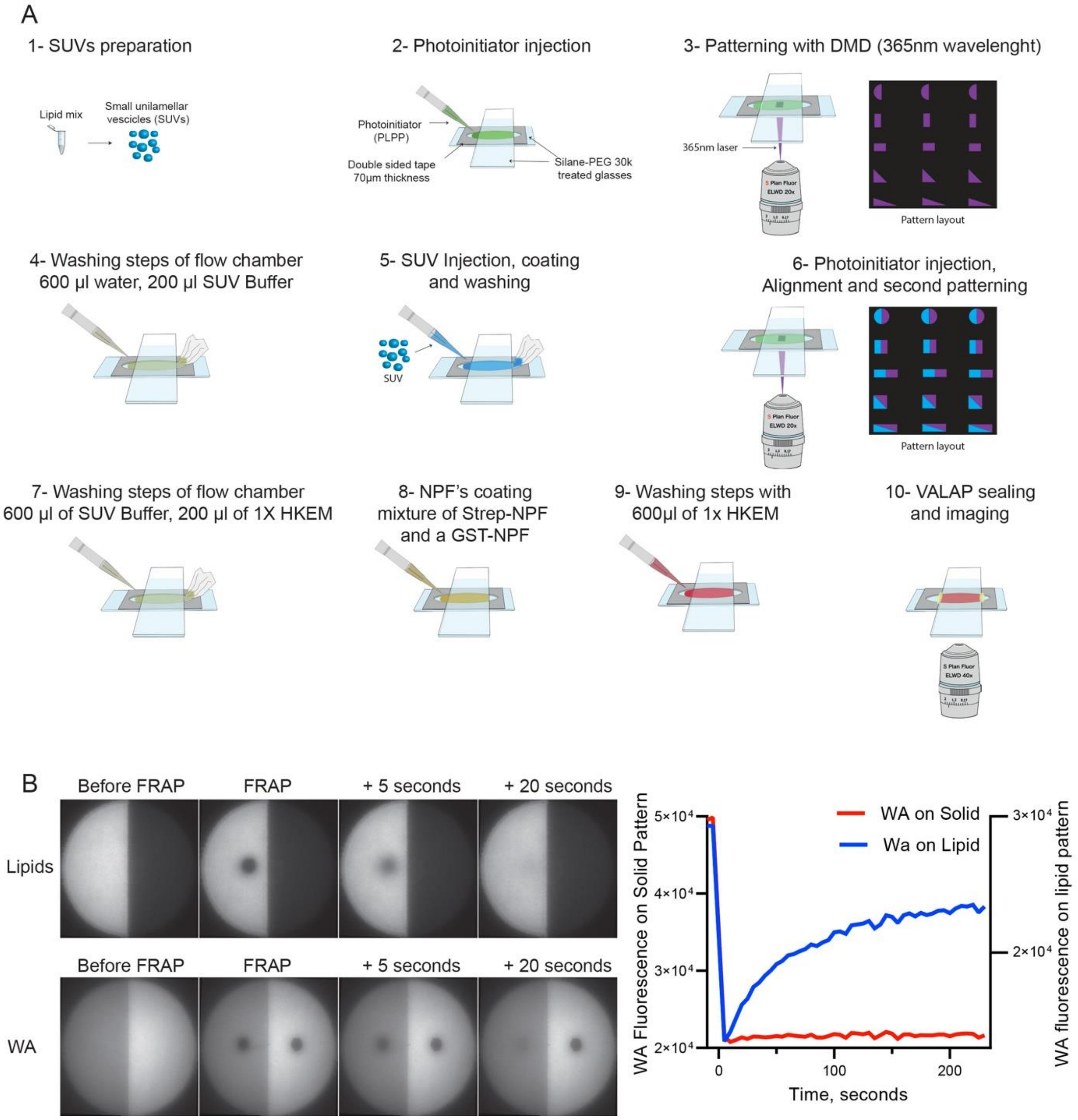


**Supporting Figure 3. Heterogeneous micropatterns imposing two distinct frictions to acto-myosin networks.**

**A.** Method to prepare heterogeneous micropatterns. **B.** Evaluation of diffusivity of lipids and NPF (WA) on heterogeneous patterns.

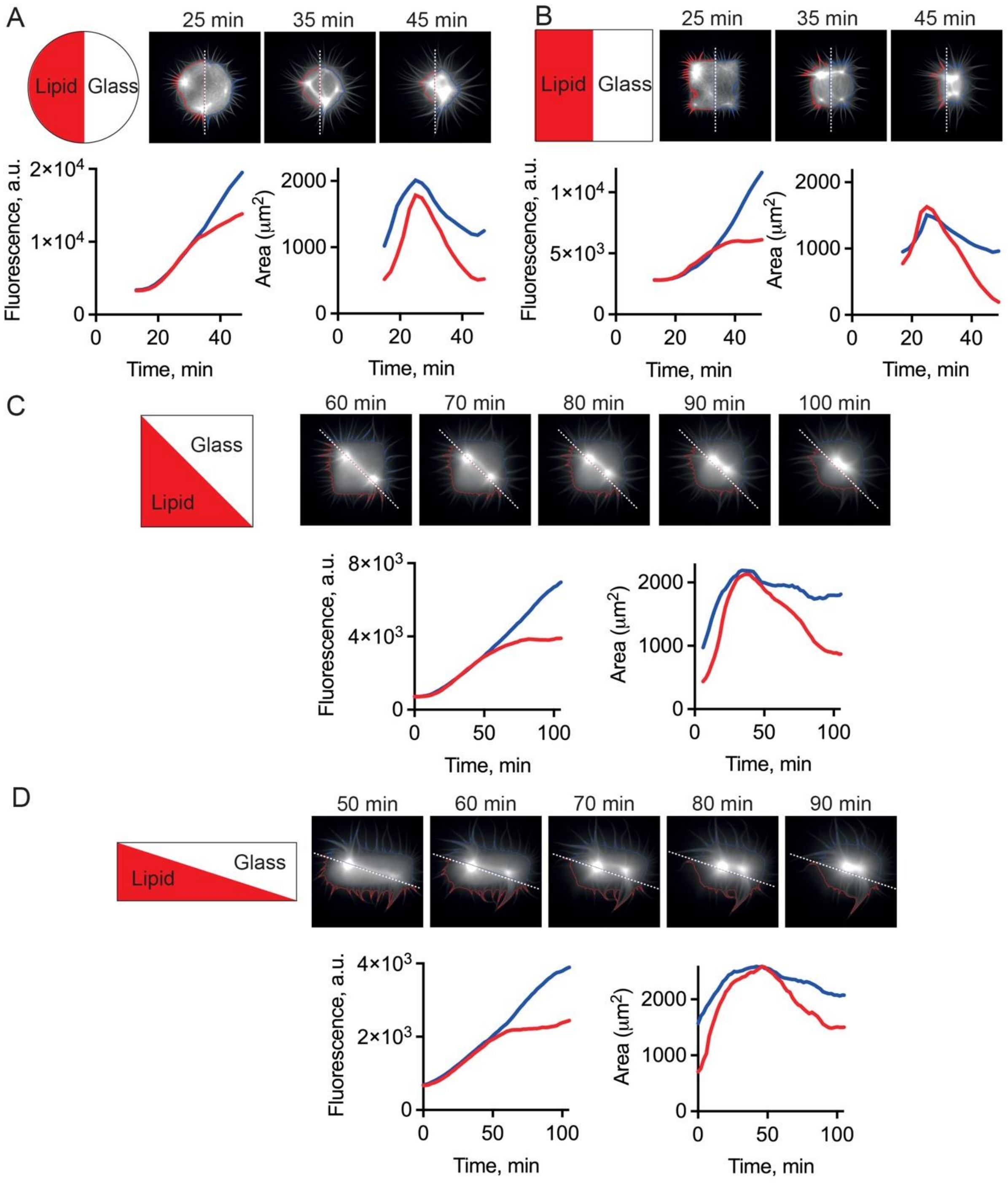


**Supporting Figure 4. Kinetics of actin polymerization and contraction on heterogeneous micropatterns.**

Top: examples snapshots of contraction on heterogeneous pattern (disc **(A)**, square **(B)**, asymmetric square **(C)**, asymmetric rectangle **(D)**). Actin network grown on lipids or glass is contoured with red or blue line respectively. Dotted white line represents the limit between the two substrates. Bottom left: Intensity of actin network grown on lipids (red) or glass (blue) for the images shown above. Bottom right: Area of actin network grown on lipids (red) or glass (blue) for the images shown above.

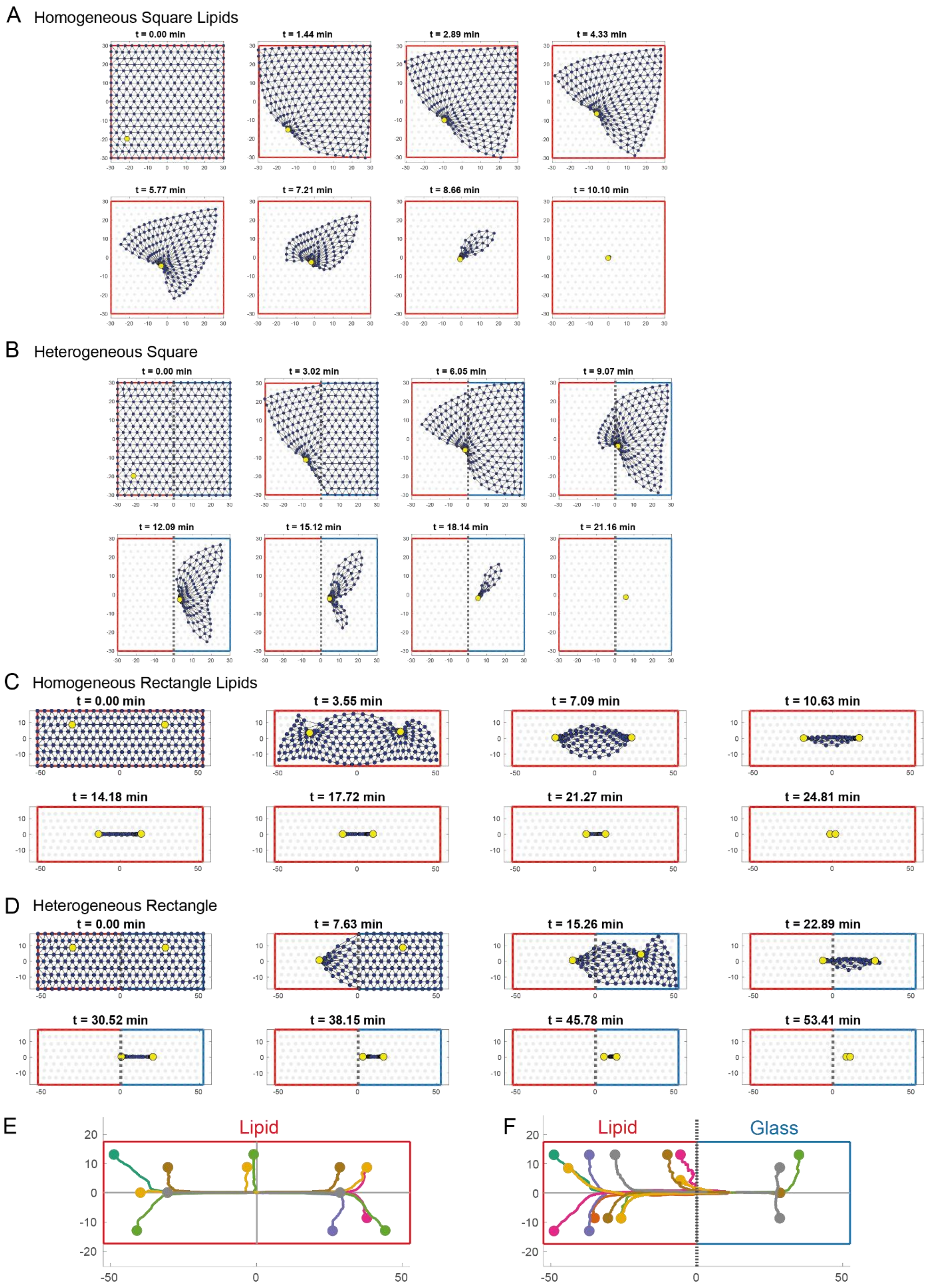


**Supporting Figure 5. Additional simulations with distinct myosin distribution.**

**A.** Example of deformation map predicted by the model for an actin network polymerized on a square lipid micropattern (asymmetric initial myosin distribution). **B.** Example of deformation map predicted by the model for an actin network polymerized on a heterogeneous square micropattern (asymmetric initial myosin distribution). **C.** Example of deformation map predicted by the model for an actin network polymerized on a rectangular lipid micropattern (symmetric initial myosin distribution). **D.** Same as (C) on an heterogenous glass/lipid pattern. **E, F.** Predictions of myosin trajectories for an actin network contracting on a homogeneous (E) or heterogeneous (F) rectangular micropattern.

#### Movies Legends

**Movie 1 (related to Figure 1B). Branched actin assembly on lipid micropattern**
TIRF Imaging of branched actin network assembly on lipid micropattern. Data is also shown in Figure 1B. Area of the zoom is 41μm². 1 image was taken every 5 seconds during 30 minutes. The movie was compressed in JPEG at 45 frames per seconds.

**Movie 2 (related to Figure 1C). Characterization of the diffusion property of the lipid micropattern.**
TIRF imaging before and after FRAP (zone diameter: 10μm) on lipids, NPF (WA) and Actin (after network polymerization). Data is also shown in Figure 1C. Movie duration is 25 seconds. Movie playback is 25 frames per second.

**Movie 3 (related to Figure 1D). Comparison of the efficiency of actin network growth on lipid versus glass micropatterns.**
TIRF imaging of branched actin assembly on lipid and glass disc micropatterns. Data is also shown in Figure 1D. 1 image was taken every 20 seconds during 40 minutes. The movie was compressed in JPEG at 20 frames per second.

**Movie 4 (related to Figure 2A). Examples of contraction of disk-shaped actin networks on glass or lipid micropatterns.** 4 examples of TIRF imaging of actin network contraction on a glass or lipid disc (diameter 68 μm) micropattern. 1 image was taken every 2 minutes during 45 minutes. The movie was compressed in JPEG at 10 frames per second.

**Movie 5 (related to Figure 2B). Examples of contraction of square-shaped actin networks on glass or lipid micropatterns.** Four examples of TIRF imaging of actin network contraction on a glass or lipid square (length 60 μm) micropattern. 1 image was taken every 2 minutes during 45 minutes. The movie was compressed in JPEG at 10 frames per second.

**Movie 6 (related to Figure 3D, Glass). Examples of contraction of square-shaped actin networks on glass micropatterns.**
Four examples of TIRF imaging of actin network contraction on a glass square (length 60 μm) micropattern. Actin is in red and myosin VI in green. The borders and the medians of the square are drawn in yellow. 1 image was taken every minute during 100 minutes. The movie was compressed in JPEG at 15 frames per second.

**Movie 7 (related to Figure 3D, Lipids). Examples of contraction of square-shaped actin networks on lipid micropatterns.**
Four examples of TIRF imaging of actin network contraction on a lipid square (length 60 μm) micropattern. Actin is in red and myosin VI in green. The borders and the medians of the square are drawn in yellow. 1 image was taken every minute during 30 minutes. The movie was compressed in JPEG at 5 frames per second.

**Movie 8. Examples of contraction of full-disc shaped actin networks on lipid micropatterns.**
Eight examples of time-lapse imaging of actin network contraction on a lipid full disc. Actin is in red, myosin VI is in green and lipid pattern is in blue. 1 image was taken every 2 minutes for 80 minutes. The movie was compressed in JPEG at 10 frames per second.

**Movie 9. Examples of contraction of disc shaped actin networks on heterogeneous micropatterns.**
Eight examples of time-lapse imaging of actin network contraction on a heterogeneous disc. Actin is in red, myosin VI is in green and lipid pattern is in blue. 1 image was taken every 2 minutes for 96 minutes. The movie was compressed in JPEG at 10 frames per second.

**Movie 10. Examples of contraction of full-square shaped actin networks on lipid micropatterns.**
Eight examples of time-lapse imaging of actin network contraction on a lipid full square. Actin is in red, myosin VI is in green and lipid pattern is in blue. 1 image was taken every 2 minutes. The movie was compressed in JPEG at 10 frames per second.

**Movie 11. Examples of contraction of square shaped actin networks on heterogeneous micropatterns.**
Eight examples of time-lapse imaging of actin network contraction on a heterogeneous square. Actin is in red, myosin VI is in green and lipid pattern is in blue. 1 image was taken every 2 minutes for 96 minutes. The movie was compressed in JPEG at 10 frames per second.

**Movie 12. Examples of contraction of square shaped actin networks on diagonal heterogeneous micropatterns.**
Eight examples of time-lapse imaging of actin network contraction on a diagonal heterogeneous square. Actin is in red, myosin VI is in green and lipid pattern is in blue. 1 image was taken every 2 minutes for 138 minutes. The movie was compressed in JPEG at 10 frames per second.

**Movie 13. Examples of contraction of full-rectangle shaped actin networks on lipid micropatterns.**
Four examples of time-lapse imaging of actin network contraction on a lipid full rectangle. Actin is in red, myosin VI is in green and lipid pattern is in blue. 1 image was taken every 2 minutes for 138 minutes. The movie was compressed in JPEG at 10 frames per second.

**Movie 14. Examples of contraction of rectangle shaped actin networks on heterogeneous micropatterns.**
Eight examples of time-lapse imaging of actin network contraction on a heterogeneous rectangle. Actin is in red, myosin VI is in green and lipid pattern is in blue. 1 image was taken every 2 minutes for 114 minutes. The movie was compressed in JPEG at 10 frames per second.

**Movie 15. Examples of contraction of rectangle shaped actin networks on diagonal heterogeneous micropatterns.**
Eight examples of time-lapse imaging of actin network contraction on a diagonal heterogeneous rectangle. Actin is in red, myosin VI is in green and lipid pattern is in blue. 1 image was taken every 2 minutes for 170 minutes. The movie was compressed in JPEG at 10 frames per second.

**Movie 16. Simulation of the contraction of square shaped actin network on lipid micropattern with a symmetric initial myosin distribution**
Simulation of actin network contraction on a lipid square micropattern with a symmetric initial myosin distribution. Myosin foci are shown in yellow and contour of the pattern in red. The movie is played at 45 frames per seconds.

**Movie 17. Simulation of the contraction of square shaped actin network on a heterogeneous micropattern with a symmetric initial myosin distribution**
Simulation of actin network contraction on a heterogeneous square micropattern with a symmetric initial myosin distribution. Myosin foci are shown in yellow, lipid part of the pattern in red, glass part of the pattern in blue. The movie is played at 85 frames per seconds.

**Movie 18. Simulation of the contraction of square shaped actin network on lipid micropattern with an asymmetric initial myosin distribution**
Simulation of actin network contraction on a lipid square micropattern with an asymmetric initial myosin distribution. Myosin foci are shown in yellow and contour of the pattern in red. The movie is played at 20 frames per seconds.

**Movie 19. Simulation of the contraction of square shaped actin network on a heterogeneous micropattern with an asymmetric initial myosin distribution**
Simulation of actin network contraction on a heterogeneous square micropattern with an asymmetric initial myosin distribution. Myosin foci are shown in yellow, lipid part of the pattern in red, glass part of the pattern in blue. The movie is played at 40 frames per seconds.
